# Meta-analytic evidence for distinct neural correlates of conditioned vs. verbally induced placebo analgesia

**DOI:** 10.1101/2025.05.21.655287

**Authors:** Tamas Spisak, Helena Hartmann, Matthias Zunhammer, Balint Kincses, Katja Wiech, Tor D. Wager, Ulrike Bingel, the Placebo Imaging Consortium

## Abstract

Placebo analgesia demonstrates that belief and expectation can significantly alter pain, even without active treatment. Placebo analgesia can be induced through verbal suggestion, classical conditioning, or their combination, though the role of conditioned neural responses above and beyond effects of verbal instructions remains unclear. We conducted a systematic meta-analysis of individual participant data from 16 within-participant placebo neuroimaging studies (*n* = 409), employing univariate and multivariate analyses to identify shared and distinct mechanisms of placebo analgesia induced by suggestions alone versus suggestions combined with conditioning. Both techniques increased activity during pain in the dorsolateral prefrontal and inferior parietal cortices and decreased activation in the insula, putamen, and primary sensory areas. Adding conditioning enhanced engagement of regions associated with context representation and pain modulation (e.g., dorsolateral/dorsomedial prefrontal cortices) and decreases in nociceptive regions (e.g., primary sensory and insular areas). Conditioning also strengthened the association between analgesia and nociceptive activity, as reflected in the Neurologic Pain Signature. Combining conditioning with instructions yielded greater analgesia, mediated by increased ventromedial prefrontal and dorsal caudate activity, alongside decreased sensory-nociceptive and cerebellar activity. These findings suggest the two strategies rely on partially distinct mechanisms, which could be combined to optimize placebo analgesia’s clinical application.

## Introduction

Placebo effects play a significant role in health and treatment outcomes in both experimental and clinical research. Understanding the underlying mechanisms of these effects is crucial for optimizing drug development and clinical care. Placebo analgesia (PA), the relief of pain by sham treatments - like inert pills, creams, and injections - is the best-studied example of placebo effects. Placebo analgesia is typically induced through (a) verbal suggestions from an experimenter or physician that promote positive expectations, and (b) classical conditioning techniques that offer direct experiences of pain relief by experimentally modifying nociceptive input ^1,2^. Most commonly, a combination of these two methods is employed, which has been shown to enhance the effectiveness of placebo treatments ^1,3^.

Extensive neuroscientific research over the past few decades has demonstrated that placebo analgesia (PA) is a complex biopsychosocial phenomenon, involving distinct central nervous system mechanisms that affect pain perception ^2,4^. Subjective changes in pain perception have been associated with altered processing of pain and the involvement of top-down pain modulatory activity including the release of endogenous opioids ^5,6^ and other neurotransmitters (e.g., dopamine; ^7^). More specifically, PA has been linked to the engagement of the dorsolateral and ventromedial prefrontal cortices (DLPFC and VMPFC), anterior cingulate cortex (ACC) and subcortical structures such as the hypothalamus, amygdala and the periaqueductal grey (PAG) ^8–10^. Spinal cord imaging studies suggest that the influence of this top-down modulation can even extend to the spinal level, leading to observable alterations in nociceptive processing ^11,12^. Our two recent systematic individual participant level meta-analyses of 20 independent neuroimaging studies found placebo-related activity increases consistent with modulatory activity in fronto-parietal regions, and decreases in the insula, thalamus and cerebellum. However, the effect on the Neurologic Pain Signature (NPS), an established brain signature of pain processing, was negligible in size ^13–15^. These results therefore suggest that multiple pathways and mechanisms contribute to PA and largely bypass classical ascending nociceptive pathways.

At present, the question of whether verbal instruction and conditioning, as the two main methods of inducing PA, rely on the same neural pathways remains unanswered. Involvement of the descending pain modulatory system, opioid release and altered spinal cord processing have so far only been documented in studies with a conditioning component ^5,6,12,16^. Whether this is the result of selective reporting and publication bias, or whether it actually reflects different mechanisms of conditioned and verbally induced PA, remains to be investigated. Evidence from studies that consider the impact of these two induction methods on dynamic learning in the context of fear, reward, and pain suggests that the neural underpinnings of these processes may indeed differ, and that they can interact when combined, such that instructions shape brain responses during learning, particularly in the striatum and ventromedial prefrontal cortex ^17–21^. Behavioral studies indicate that conditioning enhances instruction-based placebo effects on pain ^3,22^ and event-related potentials ^23^ and creates placebo effects that are more resistant to later extinction and suggestions ^24,25^. However, to date, neuroimaging studies of placebo analgesia have not directly compared the effects of instructions alone and instructions paired with classical conditioning, in part because of the large sample sizes required to do so with sufficient power. Meta-analyses provide a valuable approach to fill this gap, as they offer a quantitative synthesis and comparison of effect sizes across multiple studies, thereby enhancing power and generalizability of findings across studies.

Here, we perform an individual participant data meta-analysis (‘mega-analysis’), to disentangle the neural mechanisms of placebo induced by instructions alone or in combination with conditioning. We analyzed person-level data from 16 within-participant functional neuroimaging studies collected as part of the [Placebo Imaging Consortium](https://placebo-imaging-consortium.github.io/) ^15^ which included behavioral and brain responses during evoked pain with and without placebo treatment (within-participant design). In these studies, PA was induced either by verbal instructions alone (INST) or by verbal instructions combined with conditioning (COND-INST).

First, we tested the hypothesis that combining instructions with conditioning (as compared to instructions only) indeed results in stronger behavioral PA. Second, we identified the brain regions engaged by both induction strategies and those showing differential placebo-related activity with conditioning. Third, we performed a mediation analysis to test whether differential activity in these regions mediates the effect of induction type on PA. Finally, we investigate differences between induction types and two established and validated brain signatures of pain, the Neurologic Pain Signature reflecting stimulus input dependent pain (NPS; ^13^) and the Stimulus Intensity Independent Pain Signature (SIIPS; ^26^).

## Results

We analyzed participant-level data from a subset of studies included in previous meta-analyses of placebo effects on the Neurologic Pain Signature (NPS) ^14^ and across the entire brain ^15^. Here, we included only within-participant studies, where person-level functional magnetic resonance imaging (fMRI) maps and pain ratings were available in both the placebo and the control conditions, allowing us to account for the strength of pain experience in the control condition and the magnitude of PA at the individual-person level. This resulted in a dataset with *k* = 16 studies and *n* = 415 participants (*n* = 409 after excluding participants with missing pain ratings). The included studies induced PA either by verbal instructions alone (INST; *k* = 5, *n* = 147) or by a combination of verbal instructions and conditioning (COND+INST; *k* = 11, *n* = 268, see Table 1 and **Supplementary Figures S1-S3**). A detailed description of the data collection procedures can be found in ^14^. In brief, we performed a systematic literature search to identify experimental fMRI studies of PA. Eligible studies met the following criteria: (a) published in English in a peer-reviewed journal; (b) constituted an empirical investigation; (c) involved human participants; (d) employed functional neuroimaging of the brain during evoked pain; and (e) induced pain under matched placebo and control conditions. The authors of all identified studies were contacted and invited to share their data. We collected single-participant, first-level, whole-brain standard space summary images of pain response (statistical parametric maps) from the original analyses, corresponding pain ratings separately for placebo and control conditions, experimental design parameters, and demographic data.

**Table 1.** Summary of included within-participant placebo neuroimaging studies.

| Study | Sample size ($n$) | Mean age (years) | Sex (% male) | Pain stimulus | Induction type | Treatment |
| --- | --- | --- | --- | --- | --- | --- |
| Atlas et al. (2012) | 21 | 25 | 48 | Heat | Verbal | IV-infusion |
| Bingel et al. (2006) | 19 | 24 | 79 | Laser | Verbal + Cond | Topical cream |
| Bingel et al. (2011) | 22 | 28 | 68 | Heat | Verbal + Cond | IV-infusion |
| Choi et al. (2011) | 15 | 25 | 100 | Electro | Verbal + Cond | IV-infusion |
| Eippert et al. (2009) | 40 | 26 | 100 | Heat | Verbal + Cond | Topical cream |
| Ellingsen et al. (2013) | 28 | 26 | 68 | Heat | Verbal | Nasal spray |
| Elsenbruch et al. (2012) | 36 | 26 | 42 | Distension | Verbal | IV-infusion |
| Freeman et al. (2015) | 24 | 27 | 50 | Heat | Verbal + Cond | Topical cream |
| Geuter et al. (2013) | 40 | 26 | 100 | Heat | Verbal + Cond | Topical cream |
| Kong et al. (2006) | 10 | 27 | 60 | Heat | Verbal + Cond | Sham acupuncture |
| Kong et al. (2009) | 12<sup>a</sup> | 26 | 42 | Heat | Verbal + Cond | Sham acupuncture |
| Lui et al. (2010) | 31 | 23 | 45 | Laser | Verbal + Cond | Sham TENS |
| Schenk et al. (2014) | 32 | 26 | 53 | Cap + Heat | Verbal | Topical cream |
| Theysohn et al. (2014) | 30 | 35 | 50 | Distension | Verbal | IV-infusion |
| Wrobel et al. (2014) | 38 | 26 | 58 | Heat | Verbal + Cond | Topical cream |
| Zeidan et al. (2015) | 17<sup>a</sup> | 28 | 47 | Heat | Verbal + Cond | Topical cream |

*Note.* Total $n = 415$ participants (409 after excluding individuals with missing pain ratings); Cap = capsaicin; IV = intra-venous; TENS = transcutaneous electrical nerve stimulation; Cond = conditioning. <sup>a</sup> Only placebo-treatment groups.

To address heterogeneity between studies, we harmonized the brain data (control –placebo contrast images) by converting the values to their quantile distance from zero, independently for each voxel within each study (see Methods for details). We then examined the total effect of induction type (*X*) on behavioral analgesia (*Y*), referred to as the “total effect” (*path c*, **Figure 1A**). Next, we conducted a mediation analysis (**Figure 1B**) to identify the brain mediators (*M*) underlying this association. In our mediation framework, *path a* corresponds to the effect of induction type on brain placebo responses (*X→M*), and *path b* corresponds to the correlation between placebo-related brain activity and behavioral PA (*M →Y*). We evaluated the mediation effect *a*b*, i.e. the joint effects of induction type on brain placebo response (*path a*) and brain placebo response on behavioral PA (*path b*). We included age, sex and pain ratings in the control condition as covariates and modeled studies with dummy regressors orthogonalized to induction type.

**Figure 1.**
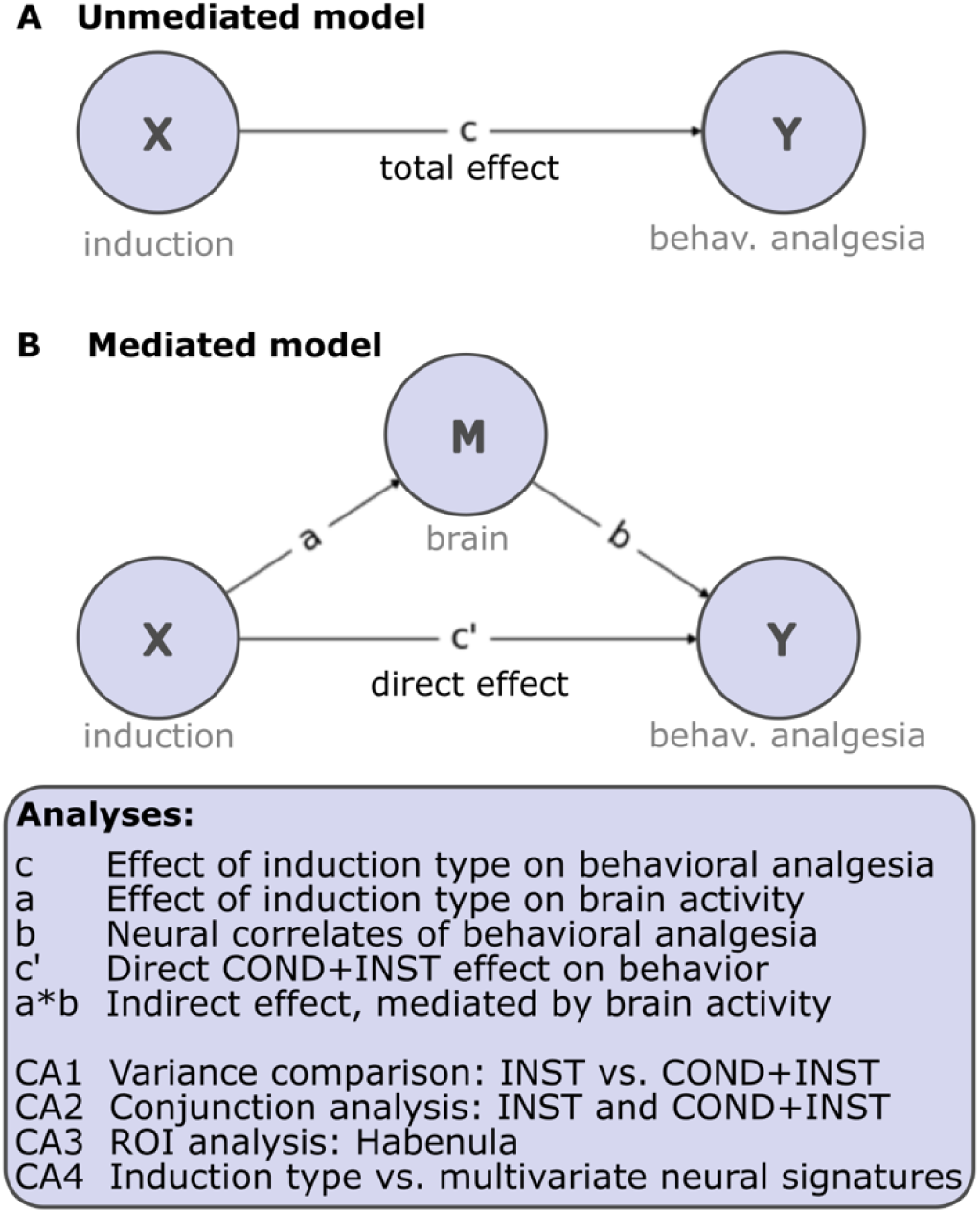
Analyses performed in the present manuscript. We fit an unmediated (A) and a mediated (B) model on placebo induction technique (**X**: INST or COND+INST), behavioral analgesia (**Y**: ctr-placebo rating difference on VAS 0-100) and the quantile-harmonized brain placebo response data (**M**: ctr-placebo contrast maps). This results in analyses quantifying the effect: **c**: induction type on behavioral analgesia, **a** induction type on brain activity, **b**: brain activity on behavioral analgesia, as well as the direct (**c’)** and indirect or mediated (**a*b)** effects. CA1: We compared the variances in the unharmonized behavioral placebo responses between INST and COND+INST; CA2: complementary conjunction analysis to identify brain regions activating in both induction types. CA3: ROI-analysis focusing on the placebo responses in the Habenula; CA4: Effect of induction type on two multivariate neural signatures: the Neurologic Pain Signature (NPS) and the Stimulus Intensity-Independent Pain Signature (SIIPS).

We performed four further analyses (CA1-4), complementary to the brain mediation analyses: (i) we compared the variances in the unharmonized placebo responses across INST and COND+INST studies (accounting for age, sex and baseline pain rating); (ii) we conducted a conjunction analysis to identify brain regions that are involved both in INST and COND+INST; (iii) we performed a confirmatory ROI-based analysis focusing on the Habenula and (iv) extracted the NPS and SIIPS scores independently for all participants and analyzed whether induction type has an effect on their association with behavioral PA.

### Path c: Behavioral results

Mean (± standard error) PA was 9.2 (± 1.83) and 12.2 (± 0.82) points on a 0-100 visual analogue scale (VAS), respectively, indicating that combining instructions with conditioning led to an additional pain reduction of 3 points, on average (as compared to verbal instructions alone, see **Figure 2**). These absolute effect size translated to a mean (± standard deviation across studies) standardized Hedges’ *g* effect of 0.367 (± 0.121) and 0.744 (± 0.278) for INST and COND+INST studies, indicating ‘small-to-moderate’ and ‘moderate-to-large’ effects, respectively. The difference 0.377 (± 0.303 was found to be statistically significant (*T*(14) = 2.86, *p* = .006, one-tailed). When corrected for age, sex, pain ratings in the control condition, and inter-study differences, the absolute effect size decreased to 1.34 VAS-points (*p* = .049, one-tailed; 95% CI: −0.247, 2.926). In the same model, we found that PA was overall strongly significant (11.15 points on the VAS pooled across induction types; *p* < .001; 95% CI: 9.62, 12.67) and it varied significantly across studies (*F*(14) = 2.44, *p* = .003). Furthermore, we found that PA was statistically significantly associated with the reported pain intensity in the control conditions (0.30 points more PA with every one-point increase in control pain; *p* < .0001, 95% CI: 0.22, 0.39) and it was weakly but statistically significantly correlated with age (0.29 points less PA with every additional year; *p* = .048; 95% CI: −0.58, −0.002). See Table 2 for details.

**Figure 2.**
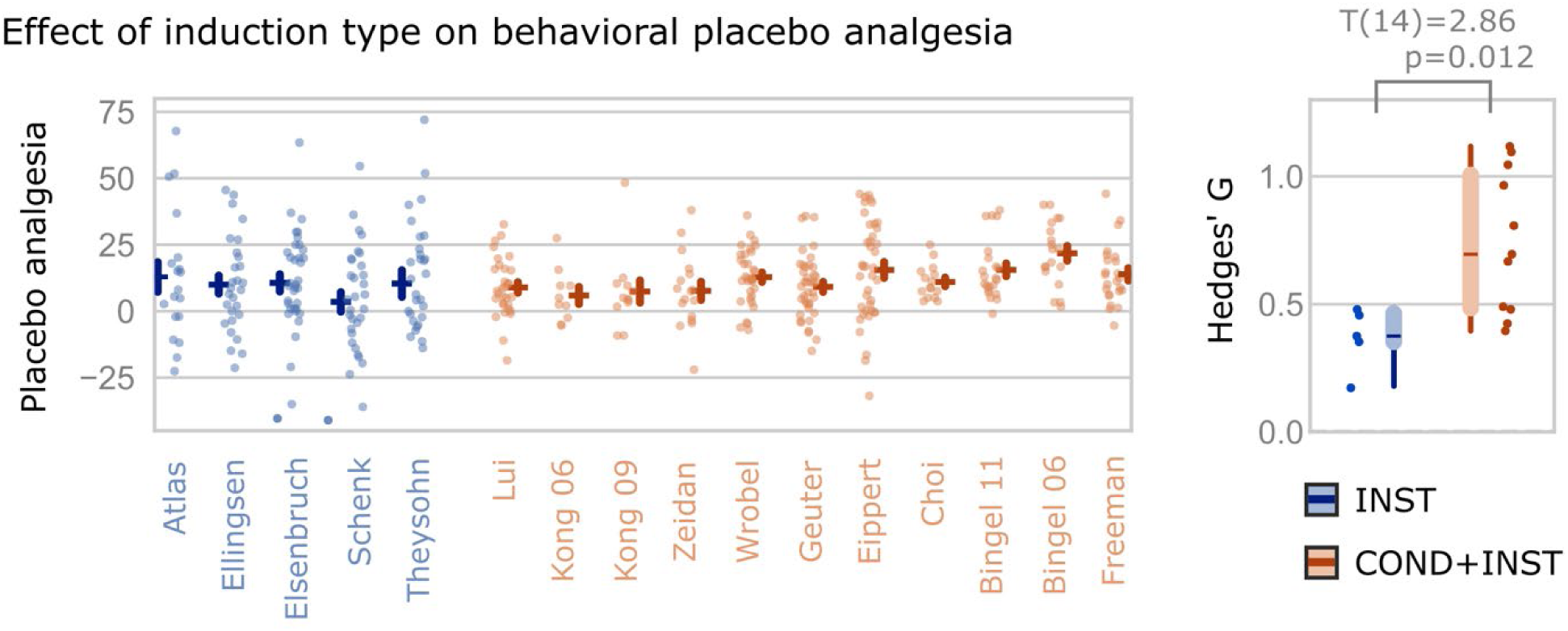
Applying conditioning together with verbal instructions results in stronger behavioral placebo effects than verbal instructions alone. Placebo analgesic effects in studies using verbal instruction alone (INST, shown in blue) and studies using a combination of verbal instructions and conditioning (COND+INST, shown in orange). On the left side, the y-axis shows the difference in the 0-100 VAS pain intensity rating between the placebo condition and the control condition. Greater values indicate a stronger behavioral placebo analgesia. Points indicate individual participants’ effects. On the right side, the standardized Hedges’ G effect sizes are shown separately for each study. Boxplots indicate the mean (and the inter-quartile and min-max ranges) of each study group. The standardized effect size difference is 0.377 (Hedges’ G) which is statistically significant with T(14) = 2.86 and p = .006, one-tailed. For results corrected for age, sex and pain rating in the control condition, see Table 2.

**Table 2.**
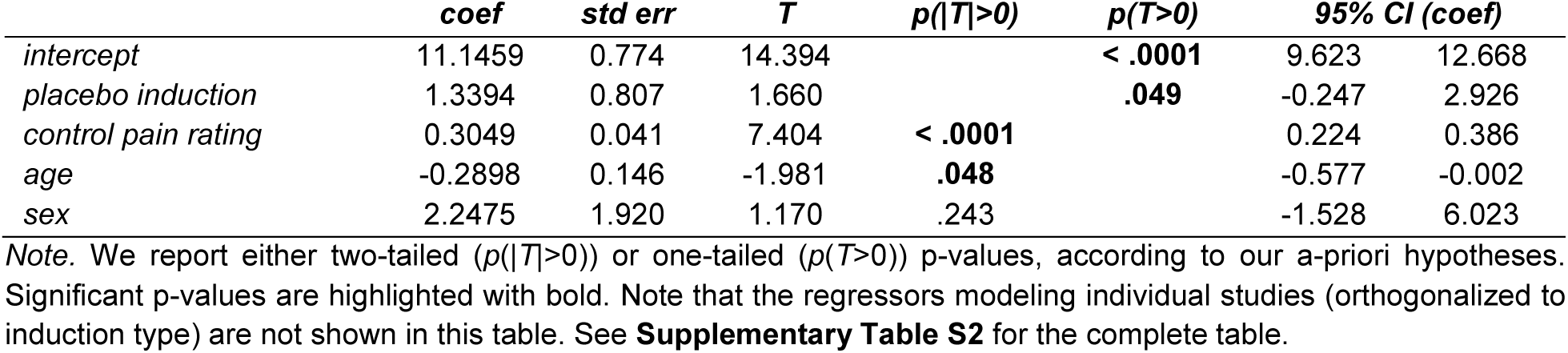
Regression coefficients, standard errors, T-scores, p-values and confidence intervals for the statistical test modelling the effect of induction type on behavioral placebo analgesia.

In our complementary analysis CA1, we tested whether the variances in the unharmonized placebo ratings are different with INST and COND+INST. First, we fit a mixed-effect model to explain the behavioral placebo effect with study as random effect and age, sex and control pain rating as fixed effects (see results in **Supplementary Table S1**). We then compared the residual variance between INST and COND+INST studies with Levene’s test and found that the variances of behavioral placebo effects with the two placebo induction types were significantly different (Levene’s test statistic: 27.2257, *p* < .0001).

### Path a: Effect of induction type on brain activity

First, we identified brain regions involved in PA both with INST and COND+INST with a statistical conjunction analysis (CA2). Our analysis revealed joint signal increases (Placebo > Control) in the DLPFC and the inferior parietal cortex (**Figure 3A**, Table 3), all surviving FDR-correction (*q* < 0.05) for multiple comparisons. Joint FDR-significant decreases (Placebo < Control) were found in the putamen, insula and the cingulate isthmus.

**Figure 3.**
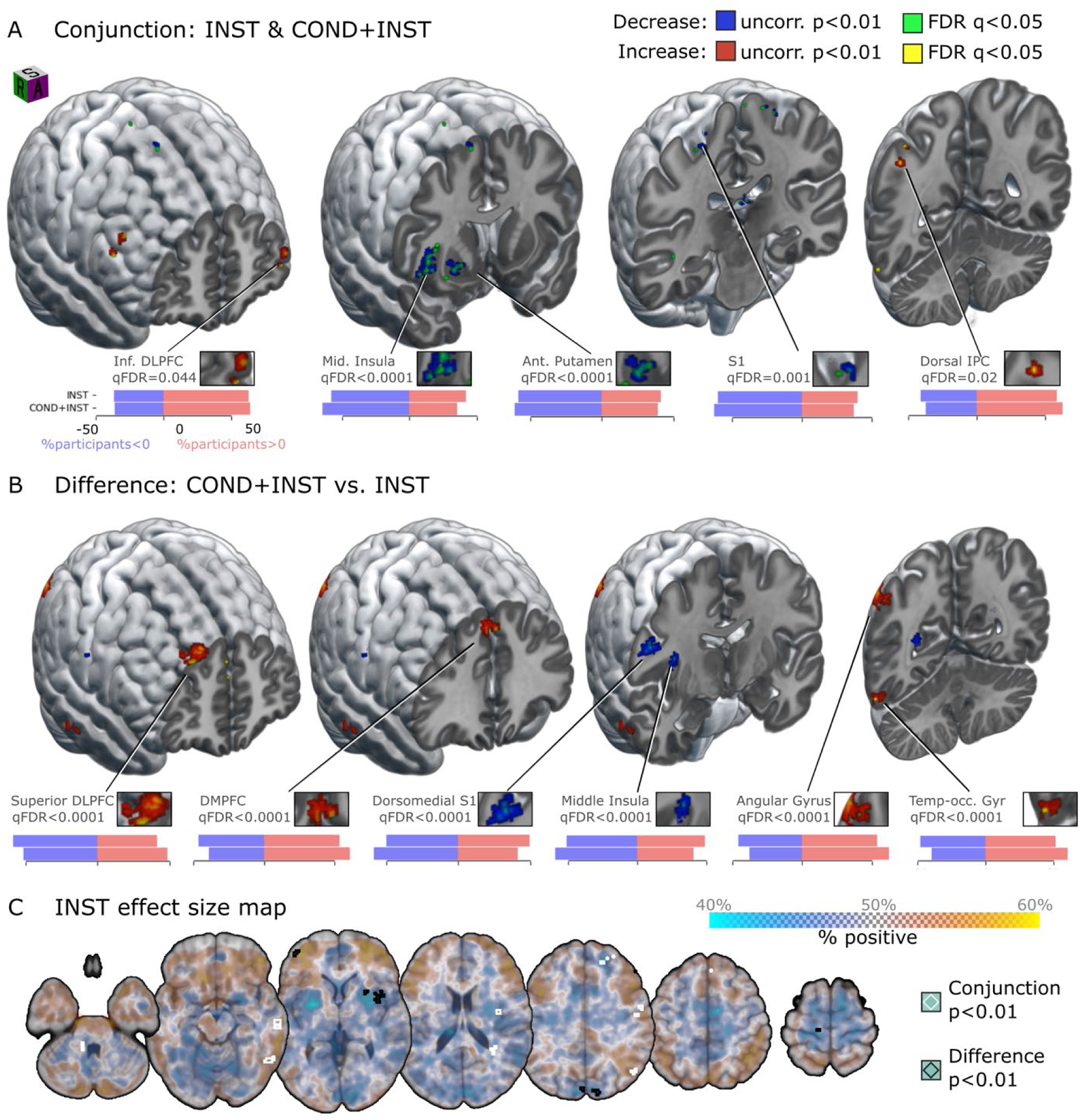
Path a: Neural correlates of placebo analgesia induction types. **(A)** Both INST and COND+INST were characterized by activations in the DLPFC and the inferior parietal cortex and deactivations in the putamen, insula and the isthmus cingulate cortex. The proportion of participants with positive (red) and negative (blue) activations in the INST and COND+INST studies is visualized by horizontal bars for selected regions. See Table 3 for a comprehensive list. **(B)** COND+INST resulted in significantly higher placebo-related activation in the middle temporal gyrus, DMPFC, DLPFC, dorsal inferior parietal cortex and somatosensory areas. As in panel A, the proportion of participants with positive (red) and negative (blue) activations in the COND and COND+INST studies is visualized by horizontal bars for selected regions. See Table 4 for a comprehensive list. **(C)** Unthresholded effect size map in the INST-studies, overlayed with the contours of the conjunction (white) and the difference (black) maps. Due to our harmonization approach, effect size translates to the ratio of participants with positive activation as shown by the colorbar. The transparency of the color map for plotting the effect size is modulated by effect size magnitude, as shown on the colorbar (smaller effects are more transparent). The spatial orientation of the three-dimensional visualizations is denoted by a cube on panel A, with letters denoting the superior (S), right (R) and anterior (A) views. In the case of uncorrected results (p < .01), we only show clusters that contain at least one voxel surviving false discovery rate correction (q < 0.05).

**Figure 4.**
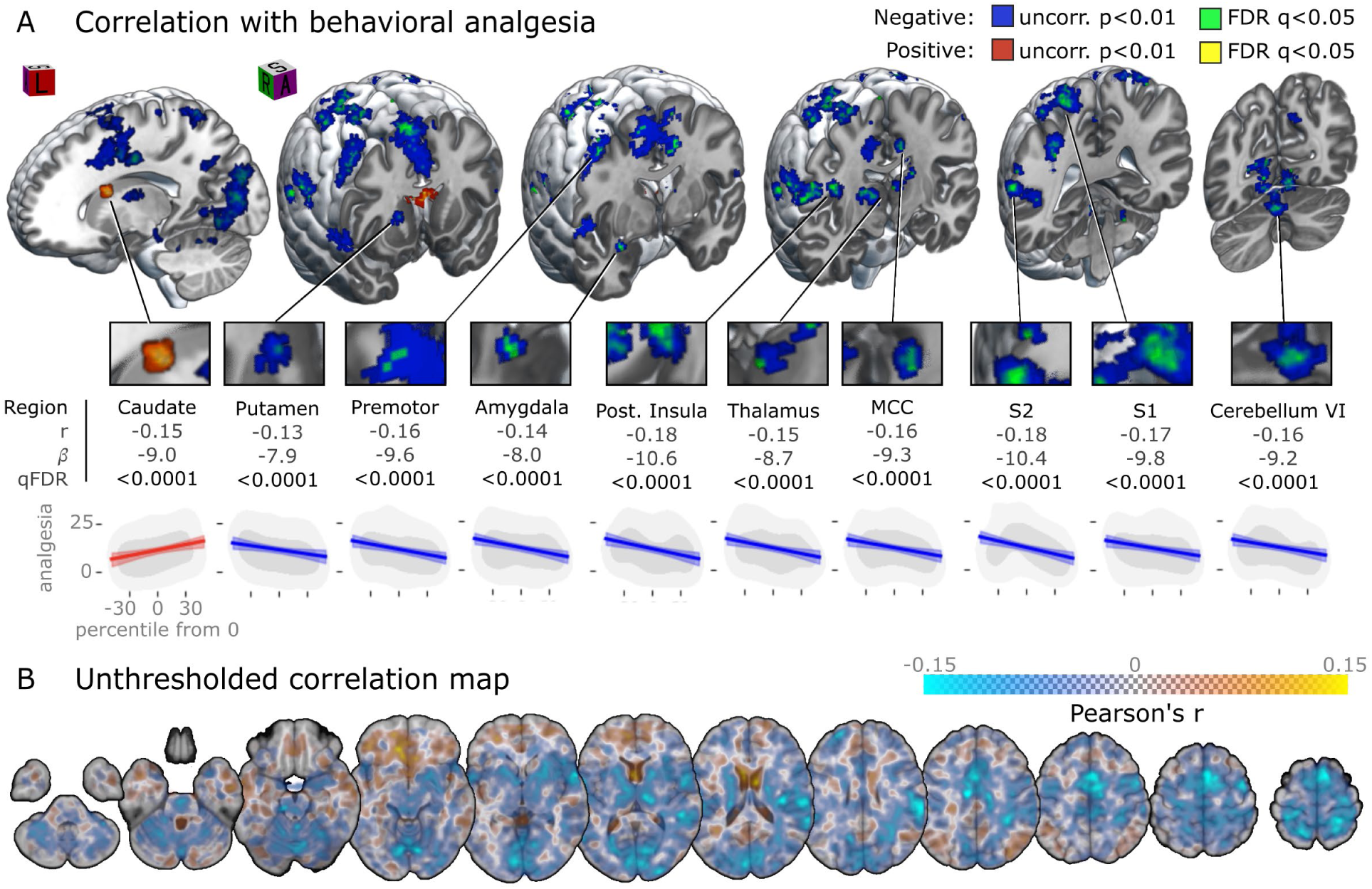
Path b: Correlation between placebo-related brain activity and behavioral placebo analgesia. **(A)** Areas where placebo-related activity reductions are significantly correlated with the degree of behavioral placebo analgesia (PA), shown in blue, include sensory-discriminative regions, mid-cingulate and premotor areas, cerebellum lobules VI and the amygdala. Only the caudate showed a significant placebo-related activity increase that positively scaled with behavioural analgesia. Correlations are depicted as regression lines (with 95% confidence intervals) overlayed on the data distribution (33 and 66% density iso-contours shown by the light and dark gray areas, respectively), for selected regions. Pearson’s correlation r, expected effect on ratings by a 10% change in population percentile (β) and FDR q-values are given for these regions. See Table 5 for a comprehensive list. **(B)** Unthresholded correlation map. The transparency of the color map for plotting the effect size is modulated by effect size magnitude, as shown on the colorbar (smaller effects are more transparent). The spatial orientation of the three-dimensional visualizations is denoted by a cube on panel A, with letters denoting the superior (S), right (R) and anterior (A) views. In the case of uncorrected results (p < .01), we only show clusters that contain at least one voxel surviving FDR correction (q < 0.05).

**Table 3.**
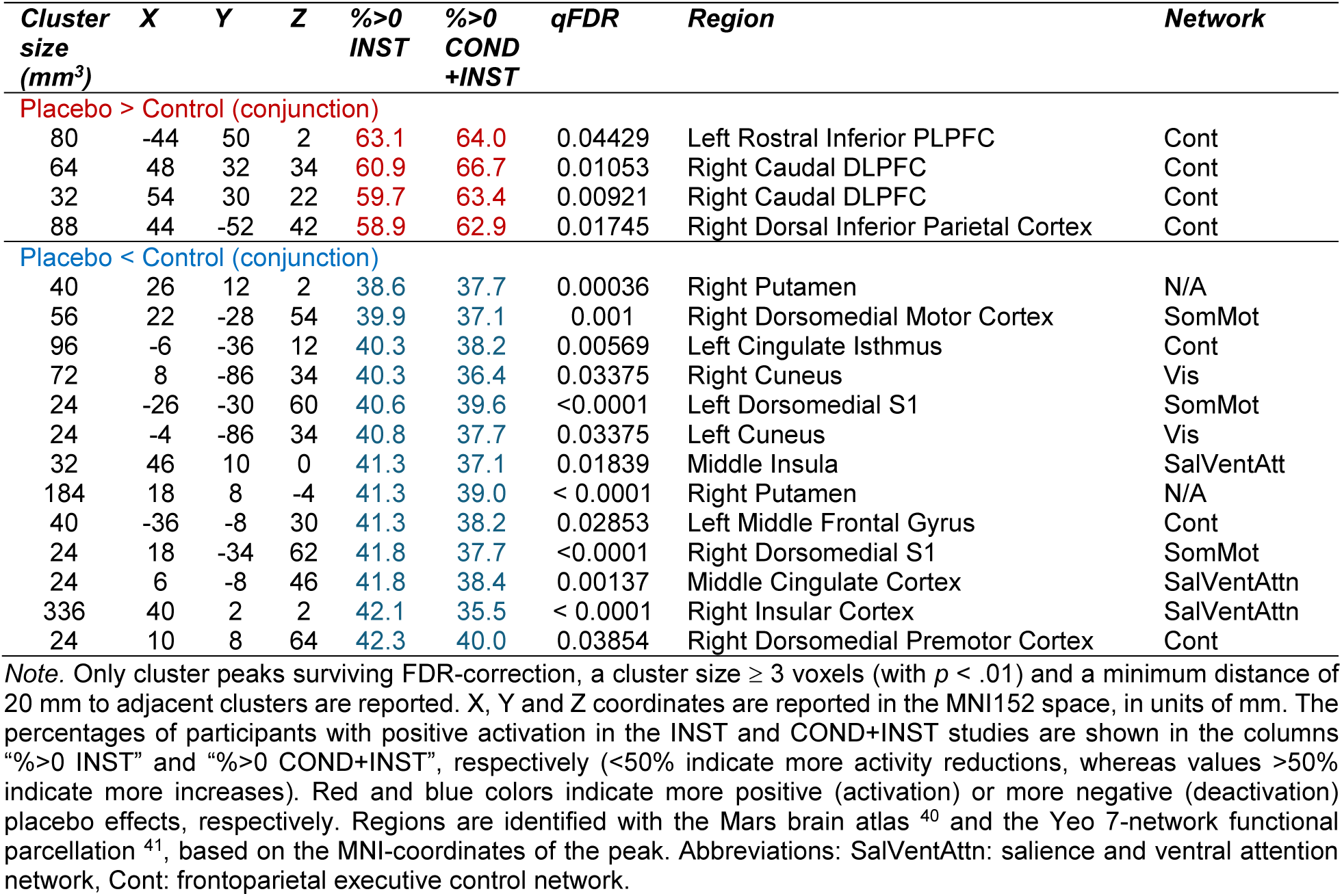
Significant activations in the INST & COND+INST conjunction analysis.

| Cluster size (mm<sup>3</sup>) | X | Y | Z | %>0 INST | %>0 COND+INST | qFDR | Region | Network |
| --- | --- | --- | --- | --- | --- | --- | --- | --- |
| **Placebo > Control (conjunction)** |  |  |  |  |  |  |  |  |
| 80 | -44 | 50 | 2 | 63.1 | 64.0 | 0.04429 | Left Rostral Inferior PLPFC | Cont |
| 64 | 48 | 32 | 34 | 60.9 | 66.7 | 0.01053 | Right Caudal DLPFC | Cont |
| 32 | 54 | 30 | 22 | 59.7 | 63.4 | 0.00921 | Right Caudal DLPFC | Cont |
| 88 | 44 | -52 | 42 | 58.9 | 62.9 | 0.01745 | Right Dorsal Inferior Parietal Cortex | Cont |
| **Placebo < Control (conjunction)** |  |  |  |  |  |  |  |  |
| 40 | 26 | 12 | 2 | 38.6 | 37.7 | 0.00036 | Right Putamen | N/A |
| 56 | 22 | -28 | 54 | 39.9 | 37.1 | 0.001 | Right Dorsomedial Motor Cortex | SomMot |
| 96 | -6 | -36 | 12 | 40.3 | 38.2 | 0.00569 | Left Cingulate Isthmus | Cont |
| 72 | 8 | -86 | 34 | 40.3 | 36.4 | 0.03375 | Right Cuneus | Vis |
| 24 | -26 | -30 | 60 | 40.6 | 39.6 | <0.0001 | Left Dorsomedial S1 | SomMot |
| 24 | -4 | -86 | 34 | 40.8 | 37.7 | 0.03375 | Left Cuneus | Vis |
| 32 | 46 | 10 | 0 | 41.3 | 37.1 | 0.01839 | Middle Insula | SalVentAtt |
| 184 | 18 | 8 | -4 | 41.3 | 39.0 | < 0.0001 | Right Putamen | N/A |
| 40 | -36 | -8 | 30 | 41.3 | 38.2 | 0.02853 | Left Middle Frontal Gyrus | Cont |
| 24 | 18 | -34 | 62 | 41.8 | 37.7 | <0.0001 | Right Dorsomedial S1 | SomMot |
| 24 | 6 | -8 | 46 | 41.8 | 38.4 | 0.00137 | Middle Cingulate Cortex | SalVentAttn |
| 336 | 40 | 2 | 2 | 42.1 | 35.5 | < 0.0001 | Right Insular Cortex | SalVentAttn |
| 24 | 10 | 8 | 64 | 42.3 | 40.0 | 0.03854 | Right Dorsomedial Premotor Cortex | Cont |

Note. Only cluster peaks surviving FDR-correction, a cluster size $\geq 3$ voxels (with $p < .01$ ) and a minimum distance of 20 mm to adjacent clusters are reported. X, Y and Z coordinates are reported in the MNI152 space, in units of mm. The percentages of participants with positive activation in the INST and COND+INST studies are shown in the columns “%>0 INST” and “%>0 COND+INST”, respectively (<50% indicate more activity reductions, whereas values >50% indicate more increases). Red and blue colors indicate more positive (activation) or more negative (deactivation) placebo effects, respectively. Regions are identified with the Mars brain atlas<sup>40</sup> and the Yeo 7-network functional parcellation<sup>41</sup>, based on the MNI-coordinates of the peak. Abbreviations: SalVentAttn: salience and ventral attention network, Cont: frontoparietal executive control network.

**Table 4.** Significant differences between placebo induction types INST and COND+INST.

| Cluster size (mm<sup>3</sup>) | X | Y | Z | %>0 INST | %>0 COND+INST | Diff. | qFDR | Region | Network |
| --- | --- | --- | --- | --- | --- | --- | --- | --- | --- |
| **COND+INST > INST** |  |  |  |  |  |  |  |  |  |
| 440 | 20 | 48 | 40 | 49.1 | 59.8 | 10.7 | < 0.0001 | Right Rostral DLPFC | Default |
|  | 20 | 50 | 42 | 53.3 | 62.8 | 9.5 | < 0.0001 | Right Rostral DLPFC | Default |
|  | 18 | 40 | 40 | 50.6 | 58.3 | 7.6 | < 0.0001 | Right Rostral DLPFC | Default |
|  | 20 | 44 | 34 | 41.3 | 48.5 | 7.2 | < 0.0001 | Right Rostral Superior DLPFC | Default |
| 344 | 44 | -70 | 44 | 50.9 | 60.3 | 9.4 | < 0.0001 | Right Superior Visual Cortex | Vis |
|  | 44 | -66 | 42 | 53.3 | 61.9 | 8.6 | < 0.0001 | Right Superior Visual Cortex | Vis |
|  | 48 | -66 | 44 | 54.0 | 62.2 | 8.2 | < 0.0001 | Right Superior Visual Cortex | Vis |
| 64 | -4 | 32 | 42 | 52.1 | 61.1 | 9.0 | < 0.0001 | Left Caudal DMPFC | Cont |
| 72 | 60 | -22 | -14 | 51.8 | 60.6 | 8.8 | < 0.0001 | Right Rostral MTG | Default |
| 32 | 50 | -62 | 48 | 51.8 | 60.5 | 8.7 | < 0.0001 | Right Dorsal Inf. Parietal Cort. | Cont |
| 72 | 56 | -58 | -16 | 51.8 | 60.4 | 8.6 | < 0.0001 | Right Lateral Visual Cortex | DorsAttn |
| 96 | 2 | 32 | 44 | 53.5 | 60.9 | 7.4 | < 0.0001 | Right Caudal DMPFC | Cont |
| 128 | 0 | 46 | 28 | 50.4 | 56.7 | 6.3 | < 0.0001 | Left Rostral Medial PFC | Default |
| 24 | 20 | 66 | 14 | 58.2 | 63.6 | 5.4 | < 0.0001 | Right Rostral Superior DLPFC | Cont |
| **COND+INST < INST** |  |  |  |  |  |  |  |  |  |
| 32 | -8 | -44 | -32 | 49.4 | 40.4 | -9.3 | < 0.0001 | Left Medial ITG | Cont |
|  | -6 | -44 | -30 | 46.7 | 37.9 | -8.8 | < 0.0001 | Left Medial ITG | Cont |
| 128 | -22 | -20 | 42 | 44.0 | 34.8 | -9.3 | 0.00773 | Left Dorsomedial Motor Cortex | Cont |
|  | -20 | -20 | 46 | 45.7 | 37.8 | -8.0 | < 0.0001 | Left Dorsomedial Motor Cortex | Cont |
| 328 | 50 | -6 | 28 | 50.1 | 41.0 | -9.1 | < 0.0001 | Right Dorsolateral M1 | SomMot |
|  | 52 | -10 | 32 | 49.9 | 41.4 | -8.5 | < 0.0001 | Right Dorsolateral S1 | SomMot |
|  | 50 | -12 | 28 | 49.9 | 41.5 | -8.4 | < 0.0001 | Right Dorsolateral S1 | SomMot |
| 32 | -6 | -44 | -30 | 46.7 | 37.9 | -8.8 | < 0.0001 | Left Medial ITG | Cont |
| 120 | 24 | -48 | 10 | 45.5 | 36.7 | -8.8 | < 0.0001 | Right Rostral Medial Vis. Cort. | Cont |
| 72 | -28 | -68 | 12 | 49.6 | 41.0 | -8.7 | < 0.0001 | Left Cuneus | Cont |
| 32 | 34 | -48 | 26 | 53.1 | 44.3 | -8.7 | < 0.0001 | Right Caudal MTG | Cont |
| 24 | -26 | -50 | 24 | 45.5 | 37.0 | -8.5 | < 0.0001 | Left Medial Parietal Cortex | N/A |
| 32 | 36 | -8 | 18 | 48.9 | 40.8 | -8.1 | < 0.0001 | Right Middle Insula | SalVentAtt |

Note. Only cluster peaks surviving whole-brain voxel-wise FDR-correction, a cluster size $\geq 3$ voxels (with $p < .01$ ) and a minimum distance of 20 mm to adjacent clusters are reported. X, Y and Z coordinates are reported in the MNI152 space, in units of mm. The percentage of participants with positive activation in the INST and COND+INST studies, and the difference, is shown in the rows “%>0 INST”, “%>0 COND+INST” and “Diff.”, respectively. The sign of the values is color coded; red: COND+INST > INST, blue COND+INST < INST. Regions are identified with the Mars brain atlas<sup>40</sup> and the Yeo 7-network functional parcellation<sup>41</sup>, based on the MNI-coordinates of the peak. Abbreviations: SomMot: somatomotor network, Cont: frontoparietal executive control network, Default: default mode network, Vis: visual network, DorsAttn: dorsal attention network.

**Table 5.** Regions where placebo-related activity is significantly correlated with behavioral placebo analgesia.

| Cluster size (mm<sup>3</sup>) | X | Y | Z | beta | r | qFDR | Region | Network |
| --- | --- | --- | --- | --- | --- | --- | --- | --- |
| **Positive Correlation** |  |  |  |  |  |  |  |  |
| 920 | 4 | 6 | 14 | 9.0 | 0.15 | < 0.0001 | Right Caudate | Cont |
| **Negative Correlation** |  |  |  |  |  |  |  |  |
| 1088 | 38 | -22 | 22 | -10.6 | -0.18 | < 0.0001 | Right Ventral Inferior Parietal Cortex | SomMot |
| 4160 | 62 | -26 | 28 | -10.4 | -0.18 | < 0.0001 | Right Ventral Inferior Parietal Cortex | SalVentAttn |
| 3184 | -14 | -54 | 70 | -9.7 | -0.17 | < 0.0001 | Left Medial Superior Parietal Cortex | DorsAttn |
|  | -32 | -36 | 68 | -9.2 | -0.16 | 0.00116 | Left Dorsolateral S1 | SomMot |
|  | -12 | -58 | 66 | -9.0 | -0.15 | 0.01304 | Left Medial Superior Parietal Cortex | DorsAttn |
|  | -36 | -38 | 62 | -6.5 | -0.11 | 0.02825 | Left Dorsolateral S1 | SomMot |
| 2368 | -60 | -28 | 42 | -10.2 | -0.17 | < 0.0001 | Left Dorsal Inferior Parietal Cortex | DorsAttn |
| 5056 | 44 | -34 | 64 | -9.8 | -0.17 | < 0.0001 | Right Dorsolateral S1 | DorsAttn |
|  | 16 | -54 | 64 | -8.8 | -0.15 | < 0.0001 | Right Medial Superior Parietal Cort. | DorsAttn |
|  | 32 | -36 | 70 | -9.6 | -0.16 | < 0.0001 | Right Dorsolateral S1 | SomMot |
| 2344 | 56 | 18 | -6 | -10.2 | -0.17 | 0.02979 | Right Rostral STG | Cont |
|  | 52 | 18 | -6 | -8.1 | -0.14 | < 0.0001 | Right Rostral STG | SalVentAttn |
|  | 50 | -4 | 4 | -7.8 | -0.13 | < 0.0001 | Right Ventral Motor Cortex | SomMot |
|  | 58 | -2 | 6 | -8.0 | -0.14 | 0.04023 | Right Ventral Motor Cortex | SomMot |
| 2024 | 64 | -40 | 18 | -9.2 | -0.16 | < 0.0001 | Right Caudal STG | DorsAttn |
|  | 48 | -38 | 6 | -8.5 | -0.14 | < 0.0001 | Right Caudal STG | Default |
| 12840 | -4 | -90 | 18 | -9.4 | -0.16 | < 0.0001 | Left Cuneus | Vis |
|  | 2 | -70 | -26 | -9.2 | -0.16 | < 0.0001 | Cerebellum VI anterior | N/A |
|  | -4 | -90 | 28 | -7.3 | -0.12 | < 0.0001 | Left Cuneus | Vis |
|  | -4 | -82 | 8 | -7.8 | -0.13 | < 0.0001 | Left Cuneus | Vis |
| 13296 | -12 | 2 | 42 | -9.3 | -0.16 | < 0.0001 | Left Dorsomedial Premotor Cortex | SomMot |
|  | 0 | 8 | 62 | -9.6 | -0.16 | < 0.0001 | Left Dorsomedial Premotor Cortex | SalVentAttn |
|  | 6 | -32 | 46 | -7.7 | -0.13 | < 0.0001 | Right Posterior Cingulate Cortex | SomMot |
|  | 8 | -4 | 50 | -8.2 | -0.14 | < 0.0001 | Right Dorsomedial Motor Cortex | SomMot |
| 800 | -18 | -14 | 16 | -8.0 | -0.14 | < 0.0001 | Left Thalamus | N/A |
| 928 | 16 | -18 | 12 | -8.7 | -0.15 | < 0.0001 | Right Thalamus | N/A |
| 456 | 30 | -96 | 8 | -8.9 | -0.15 | < 0.0001 | Right Caudal Medial Visual Cortex | Vis |
| 536 | -14 | -64 | 32 | -8.8 | -0.15 | 0.00906 | Left Medial Parietal Cortex | Cont |
| 368 | 32 | -6 | -14 | -8.0 | -0.14 | < 0.0001 | Right Putamen | N/A |
| 248 | 32 | -46 | 22 | -8.1 | -0.14 | < 0.0001 | Right Caudal MTG | N/A |
| 1936 | 38 | -28 | 44 | -8.5 | -0.14 | < 0.0001 | Right Dorsolateral S1 | DorsAttn |
|  | 32 | -46 | 34 | -8.3 | -0.14 | 0.00953 | Right Superior Parietal Cortex | DorsAttn |
|  | 18 | -52 | 64 | -8.4 | -0.14 | < 0.0001 | Right Dorsomedial S1 | DorsAttn |
| 1688 | 52 | 6 | 48 | -8.4 | -0.14 | < 0.0001 | Right Dorsolateral Motor Cortex | SalVentAttn |
| 384 | -58 | 4 | 2 | -8.0 | -0.14 | 0.00213 | Left Ventral Motor Cortex | SalVentAttn |
| 344 | 18 | 10 | 8 | -7.9 | -0.13 | < 0.0001 | Right Putamen | N/A |
| 448 | -26 | 54 | 28 | -7.9 | -0.13 | < 0.0001 | Left Rostral Dorsal Prefrontal Cortex | Default |

*Note.* Only cluster peaks surviving FDR-correction, a cluster size $\geq 20$ voxels (with $p < .01$ ) and a minimum distance of 20 mm to adjacent clusters are reported. X, Y and Z coordinates of the peak activation are reported in MNI152 space, in units of mm. Effect size (beta) can be interpreted as the predicted VAS difference between the participants with the least and most activated region, within a study. Effect size is converted to Pearson's correlation coefficient ( $r$ ; red:

Next, we directly compared activation during COND+INST-induced PA and INST-induced PA, corresponding to *path a* of the mediation analysis. COND+INST led to stronger activation in key modulatory areas associated with PA, such as DLPFC and dorsomedial prefrontal cortex (DMPFC), as well as in the temporo-occipital cortex, compared to INST (**Figure 3B**, Table 4).

In the case of DMPFC and the temporo-occipital cluster, increases were driven by activation during COND+INT, while DLPFC was rather driven by placebo-related deactivation within the INST condition (see horizontal bar plots on **Figure 3B** and **Figure 3C**). The combination of verbal instructions and conditioning (COND+INST) was associated with significant signal decreases in sensory and motor areas, all of which were driven by de-activation in the COND+INST condition (**Figure 3B** and **Figure 3C**). That is, only COND+INST produced significant decreases in S1 and mid-posterior insula. Our complementary ROI analysis (CA3) found that activity in the left habenula showed a significant placebo-related decrease with INST (harmonized effect size difference: −5.1%, *p* = .016) and a non-significant but similarly sized decrease with COND+INST (−5.3%, *p* = .083), replicating our previous findings ^15^. No statistically significant effects were observed in the right habenula. Additionally, the ROI analysis found no significant path a, path b, or mediation effects in either the left or right habenula.

### Path b: Correlation between placebo responses in the brain and behavioral analgesia

Next, we evaluated the association between placebo brain responses and behavioral analgesia, controlling for induction type (*path b* of the mediation analysis). We found widespread negative correlations in S1 and S2, posterior insular areas and MCC, premotor regions, amygdala, ventral thalamus and cerebellum (blue in **Figure 5**). These results indicate larger PA with greater placebo-induced decreases in activity, and are consistent with findings in our previously published analysis ^15^. In a supplementary analysis, we investigated whether a significant ‘induction type x analgesia’ interaction could be found in any voxels, but no voxels survived correction for multiple comparisons in this analysis. Nevertheless, we found a significant ‘induction type x analgesia’ interaction in the NPS (see below).

**Figure 5.**
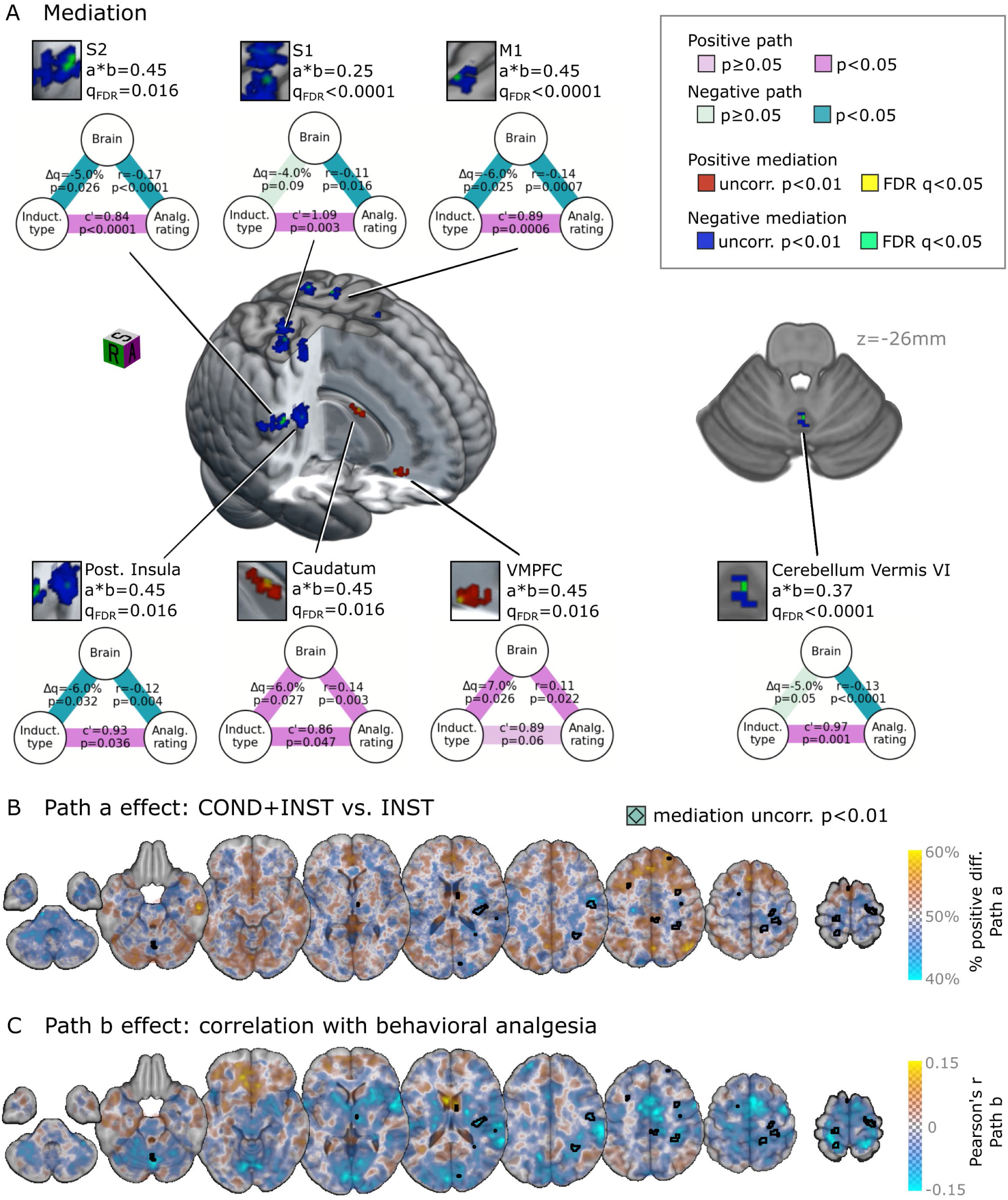
Brain mediators of placebo induction technique on behavioral placebo analgesia. **(A)** Areas where placebo induction type significantly mediates behavioral analgesia include the VMPFC (positive), caudate (positive), posterior insula (negative), S2 (negative) and S1 (negative). Red and yellow/blue and green shows regions where the effect of induction on behavioral analgesia is mediated by positive/negative, respectively, path a and path b effects. Mediation path diagrams are shown for selected regions. The path diagrams visualize the path a, path b and path c’ effects, and the corresponding uncorrected p-values. The mediation effect size (a x b) and the FDR q-value is given above the path diagrams. See Table 6 for a comprehensive list of significant regions. The spatial orientation of the three-dimensional visualizations is denoted by a cube on panel A, with S, R and A denoting the superior (gray), right (right) and anterior (anterior) views. The z-coordinate of the cerebellar slice is given in mm. See Table 6 for a comprehensive list of regions with a significant mediation effect. (**B**) Path a effect size (effect of induction type), with regions of significant mediation effect indicated with black contours. **(C)** Path b effect size (unthresholded brain-behavior correlation map), with black contours indicating significant positive mediation effect. On B and C, the transparency of the color map for plotting the effect size is modulated by the effect size magnitude, as shown on the color bars (smaller effects are more transparent). On all panels, in case of uncorrected results (p < .01), we only show clusters that contain at least one voxel surviving false discovery rate correction (q < 0.05).

**Table 6.** Regions where placebo induction type significantly mediates behavioral analgesia.

| Cluster size (mm<sup>3</sup>) | X | Y | Z | a beta | b corr | axb | qFDR | Region | Network |
| --- | --- | --- | --- | --- | --- | --- | --- | --- | --- |
| **Positive** |  |  |  |  |  |  |  |  |  |
| 40 | 8 | 2 | 16 | 6.1 | 0.14 | 0.5 | < 0.0001 | Right Caudate | N/A |
| 104 | 10 | 26 | -16 | 7.5 | 0.11 | 0.4 | 0.01642 | Right VMPFC | Cont |
| **Negative** |  |  |  |  |  |  |  |  |  |
| 880 | 54 | -18 | 24 | -5.5 | -0.17 | 0.5 | < 0.0001 | Right Dorsal Inferior Parietal Cort. | SalVentAttn |
| 176 | -16 | -46 | 68 | -6.5 | -0.16 | 0.6 | 0.00759 | Left Dorsomedial S1 | SomMot |
| 112 | 22 | -38 | 64 | -3.9 | -0.16 | 0.4 | 0.01068 | Right Dorsomedial S1 | SomMot |
| 552 | 34 | -18 | 18 | -7.3 | -0.16 | 0.6 | 0.02415 | Right Ventral Inferior Parietal Cort. | SomMot |
| 160 | 30 | -38 | 44 | -6.7 | -0.14 | 0.5 | 0.01642 | Right Superior Parietal Cortex | DorsAttn |
| 152 | -16 | -30 | 70 | -5.1 | -0.14 | 0.4 | 0.01642 | Left Dorsomedial S1 | SomMot |
| 496 | -62 | -26 | 22 | -6.3 | -0.14 | 0.5 | 0.02374 | Left Ventral Inferior Parietal Cortex | SalVentAttn |
| 200 | 26 | -20 | 68 | -6.0 | -0.14 | 0.4 | 0.03642 | Right Dorsomedial Motor Cortex | Cont |
| 48 | 2 | -60 | -26 | -5.3 | -0.13 | 0.4 | < 0.0001 | Cerebellum VI anterior | N/A |
| 64 | -14 | 0 | 62 | -5.1 | -0.13 | 0.3 | < 0.0001 | Left Dorsomedial Premotor Cortex | Cont |
| 568 | -26 | -70 | 12 | -6.8 | -0.13 | 0.5 | < 0.0001 | Left Cuneus | Cont |
| 272 | 12 | -78 | 20 | -5.7 | -0.13 | 0.4 | 0.01637 | Right Rostral Medial Visual Cortex | VisCent |
| 120 | 40 | -14 | 18 | -6.1 | -0.12 | 0.4 | 0.00125 | Right Ventral Inferior Parietal Cort. | SomMot |
| 16 | -16 | -2 | 42 | -7.2 | -0.12 | 0.5 | 0.02142 | Left Dorsomedial Premotor Cortex | Cont |
| 40 | -20 | -78 | 10 | -5.7 | -0.12 | 0.4 | 0.04147 | Left Cuneus | VisCent |
| 824 | 34 | -50 | 28 | -5.1 | -0.11 | 0.3 | < 0.0001 | Right Dorsal Inferior Parietal Cortex | Cont |
| 16 | 28 | -32 | 56 | -5.5 | -0.11 | 0.3 | < 0.0001 | Right Dorsolateral S1 | SomMot |
| 104 | 28 | -32 | 60 | -4.3 | -0.11 | 0.2 | < 0.0001 | Right Dorsolateral S1 | SomMot |
| 40 | 38 | -10 | 42 | -6.3 | -0.10 | 0.3 | < 0.0001 | Right Dorsolateral Motor Cortex | SomMot |
| 96 | -52 | -42 | 6 | -7.3 | -0.10 | 0.4 | < 0.0001 | Left Caudal STG | Default |
| 176 | 30 | -30 | 52 | -5.4 | -0.10 | 0.3 | 0.00559 | Right Dorsolateral S1 | SomMot |

*Note.* Only cluster peaks surviving FDR-correction and with a minimum distance of 20 mm to adjacent clusters are reported. X, Y and Z coordinates of the peak activation are reported in the MNI152 space, in units of mm. Path a effect size (a) can be interpreted as the predicted VAS difference between the participants with the least and most activated region. Path b (b) is given as Pearson's correlation coefficient. Effect sizes are color coded; red: positive mediator, blue: negative mediator. Regions are identified with the Mars brain atlas <sup>40</sup> and the Yeo 7-network functional parcellation <sup>41</sup>, based on the MNI-coordinates of the peak. Abbreviations: DorsAttn: dorsal attention network, Default: default mode network, SomMot: somatomotor network, Cont: frontoparietal executive control network, Vis: visual network, SalVentAttn: salience and ventral attention network,

### Brain mediators of the effect of induction type on behavioral placebo analgesia

When evaluating the mediation effect i.e. the joint effects of induction type on brain placebo response (path a) and brain placebo response on behavioral PA (path b), we found FDR-significant mediation (a*b) in several locations (**Figure 5**, Table 6). We distinguished two types of mediation effects. We refer to ‘positive mediation’ where more behavioral analgesia in COND+INST is mediated by more brain activity (i.e., both *path a* and *b* are positive). By ‘negative mediation’ we refer to cases, when more analgesia in COND+INST is mediated by less brain activity (i.e. both *path a* and *b* are negative). Placebo brain responses in the VMPFC and right caudate were found to be positive mediators of the effect of induction type on behavioral analgesia. These regions exhibit a combination of greater activation with COND+INST and a positive relationship between placebo-induced activity and analgesia. The most pronounced negative mediators were regions in the posterior insula, S1 and S2 indicating placebo analgesia being mediated by stronger deactivation of pain-related activity. These regions exhibit a combination of reduced activation with COND+INST and greater analgesia with more placebo-induced deactivation.

### Effects of conditioning on multivariate pain signatures

In the final analysis, we investigated the extent to which the placebo analgesic effect, induced by either COND-INST or INST, correlated with changes in the engagement of two established pain signatures in the brain: the Neurologic Pain Signature (NPS, **Figure 6 left**) and the Stimulus Intensity Independent Pain Signature (SIIPS, **Figure 6 right**). Pooled over induction types, the NPS exhibited a statistically significant negative correlation with behavioral analgesia (T(390)=-3.169, r=.159, p=.0015, two-tailed). When contrasting induction types, we observed a stronger NPS-analgesia association with COND+INST (T(252)=-3.025, *r* = .19, *p* =.002, two-tailed) than with INST only, (T(136)=-1.184, *r* = .1, *p* = .12, two-tailed). The difference between induction types was statistically significant (interaction: *T*(409) = −1.83, *p* = .034, two-tailed). Detailed results are presented in Table 7. The SIIPS showed a weak but significant negative correlation with behavioral analgesia both when data were pooled across the induction types (T(390)=-3.109, r=0.155, p=0.0031, two-tailed) and when analyzed separately for each (*r* = .15 and .16, *p* = .05 and .02 for INST and COND+INST, respectively), with no statistically significant interaction. For detailed results, see Table 8.

**Figure 6.**
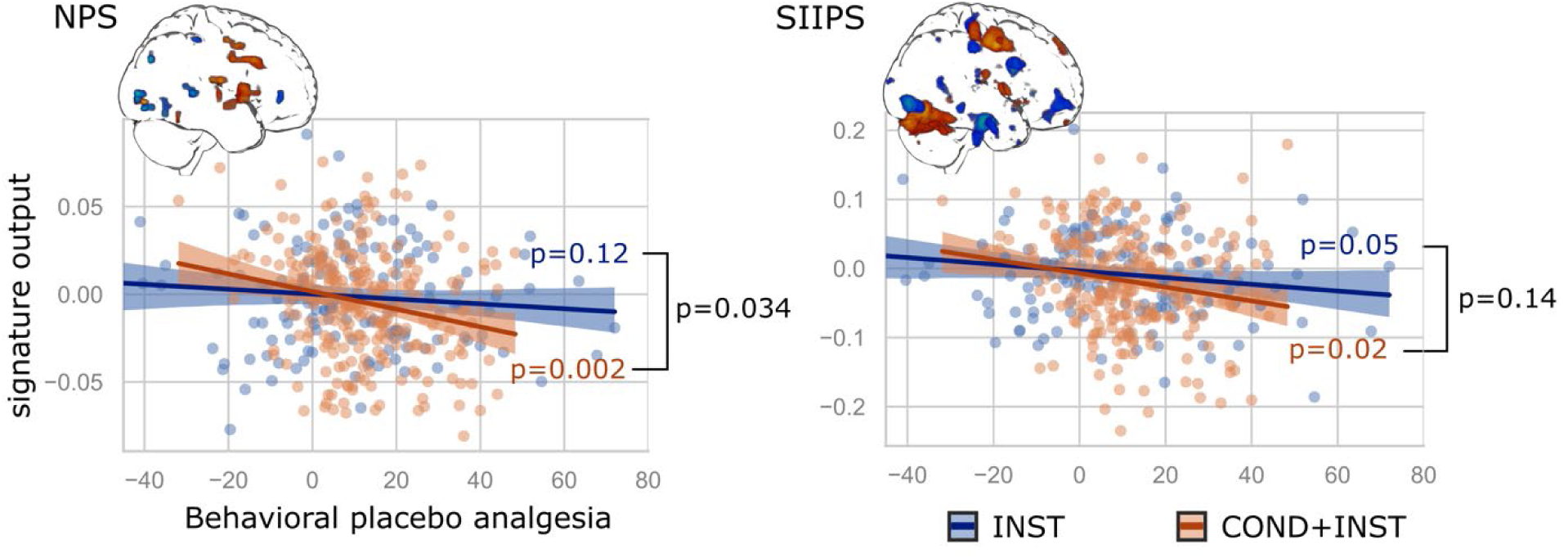
NPS, but not SIIPS, more strongly predicts behavioral analgesia induced by COND+INST than by INST alone. Scatter plots show the association between behavioral placebo analgesia and the NPS and SIIPS signature responses, respectively. Regression lines and 95% confidence intervals are plotted separately for the INST and COND+INST induction techniques. Blue and orange p-values correspond to the INST and COND+INST induction techniques, corrected for age and sex. Black p-values correspond to the interaction effect between induction technique and behavioral analgesia; with the significant p-value for NPS indicating that the association between the NPS and behavioral analgesia is significantly stronger in case of COND+INST. Glass brains represent the predictive weights of the signatures (red: positive weight, blue: negative weight).

**Table 7.** Regression coefficients, standard errors, T-scores, p-values and confidence intervals for the statistical test modelling the interaction effect between induction type and behavioral analgesia on NPS scores.

|  | **coef** | **std err** | **T** | **p** | **95% CI (coef)** |  |
| --- | --- | --- | --- | --- | --- | --- |
| *intercept* | 0.0011 | 0.002 | 0.670 | .72 | -0.0026 | 0.0047 |
| *placebo induction* | 5.17e-05 | 0.002 | 0.028 | .52 | -0.0037 | 0.00386 |
| *analgesia* | -0.0004 | 0.001 | -3.559 | **.0005** | -0.0006 | -0.00016 |
| *analgesia:induction* | -0.0002 | 8.89e-05 | -1.825 | **.034** | -0.000344 | -0.000001 |
| *control pain rating* | -6.414e-05 | 8.42e-05 | -0.762 | .21 | -0.0002 | 0.0001 |
| *age* | -0.0006 | 0.001 | -2.060 | **.014** | -0.001 | -0.0001 |
| *sex* | 0.0001 | 0.004 | 0.040 | .51 | -0.0069 | 0.0072 |

Note. Significant *p*-values are highlighted in bold. Note that the regressors modelling individual studies (orthogonalized to induction type) are not shown in this table. See **Supplementary Table S3** for the complete table.

**Table 8.**
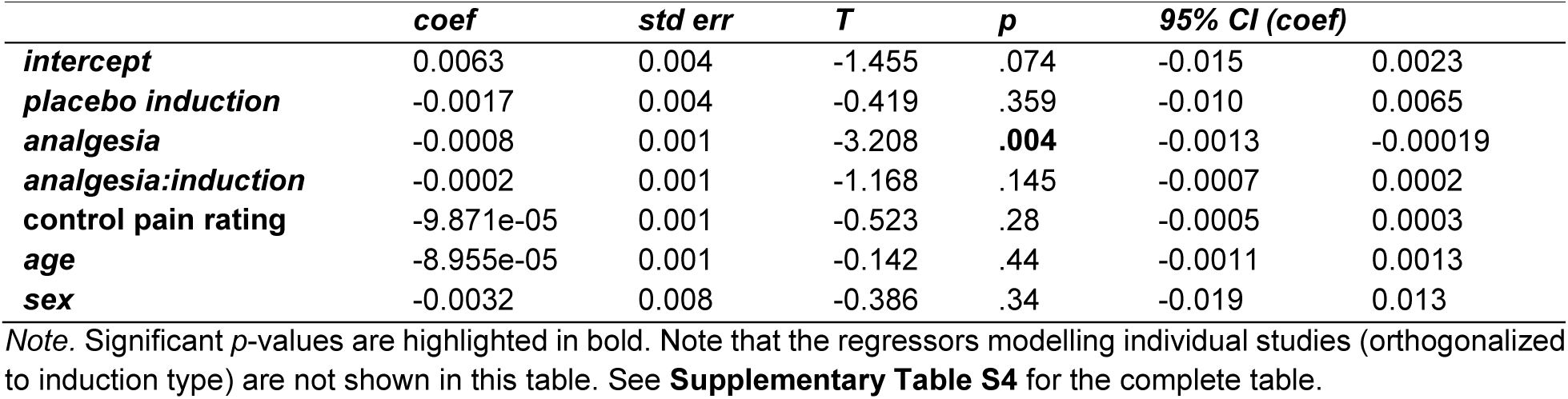
Regression coefficients, standard errors, T-scores, p-values and confidence intervals for the statistical test modelling the interaction effect between induction type and behavioral analgesia on SIIPS scores.

|  | **coef** | **std err** | **T** | **p** | **95% CI (coef)** |  |
| --- | --- | --- | --- | --- | --- | --- |
| *intercept* | 0.0063 | 0.004 | -1.455 | .074 | -0.015 | 0.0023 |
| *placebo induction* | -0.0017 | 0.004 | -0.419 | .359 | -0.010 | 0.0065 |
| *analgesia* | -0.0008 | 0.001 | -3.208 | **.004** | -0.0013 | -0.00019 |
| *analgesia:induction* | -0.0002 | 0.001 | -1.168 | .145 | -0.0007 | 0.0002 |
| *control pain rating* | -9.871e-05 | 0.001 | -0.523 | .28 | -0.0005 | 0.0003 |
| *age* | -8.955e-05 | 0.001 | -0.142 | .44 | -0.0011 | 0.0013 |
| *sex* | -0.0032 | 0.008 | -0.386 | .34 | -0.019 | 0.013 |

Note. Significant *p*-values are highlighted in bold. Note that the regressors modelling individual studies (orthogonalized to induction type) are not shown in this table. See **Supplementary Table S4** for the complete table.

## Discussion

In this systematic, individual-level meta-analysis, we examined placebo analgesia (PA) and its neural correlates when induced in healthy volunteers through either a combination of conditioning and verbal instructions (COND+INST) or verbal instructions alone (INST).

First, we confirmed that applying conditioning together with verbal instructions results in stronger PA. Second, we identified regions involved in the placebo condition regardless of induction technique, and regions where the placebo response significantly differs between induction techniques. Third, we corroborated and complemented previous results about the association between placebo response in the brain and behavioral PA, both on a voxel-wise basis and as reflected by two multivariate brain signatures. Finally, we identified brain mediators of the effect of different induction techniques on PA.

Adding conditioning to verbal instructions increased the PA effect by 33% and doubled the standardized effect size (*g* = 0.74 vs. 0.37). This is consistent with a previous meta-analysis showing a stronger behavioral effect of conditioned PA (^3^ comparison of 14 studies with sample sizes between 24 and 136 each; Cohen’s *d* = 0.85 for INST (14 studies) and *d* = 1.45 COND+INST (three studies)) and subsequent studies comparing the effect of verbal instruction with the combined use of conditioning and verbal instruction.

At the neural level, placebo analgesia induced by both INST and COND+INST was characterized by deactivation of regions involved in nociceptive processing, such as the insula, putamen and posterior cingulate cortex. Both induction techniques also led to activation in the dorsal inferior parietal cortex (IPC) and in the inferior rostral part of the DLPFC, an area that is a key component of the descending pain modulatory network ^42^ and has been implicated in orchestrating top-down pain modulation in several studies investigating PA ^8,43,44^.

By contrast, the combination of conditioning and instructions also recruited the DMPFC as a second top-down modulatory region. Further clusters were identified in the angular gyrus in the parietal cortex and superior parts of the DLPFC, which seemed to be caused by the absence of deactivation, rather than active engagement (**Figure 3B**). The recruitment of these additional descending pain modulatory regions with COND+INST was accompanied with stronger decrease in key sensory brain regions with this induction technique.

While both induction types clearly involve learning, (see ^45^), one obvious difference is the felt experience of pain reduction that is the hallmark of the conditioning procedure in a placebo manipulation. In the context of placebo analgesia, this typically involves a covert reduction in stimulation intensity, which is intended to simulate the pain relief expected from an analgesic treatment. In contrast, verbal instructions only suggest that the perceived pain intensity will be lower, without providing this first-hand experience of reduced pain. It could therefore be argued that conditioning provides a more accurate idea of pain reduction than can be conveyed by verbal instructions alone. The effect of this difference can be understood through conceptualizing perception as inference process, which have become a prominent perspective on perception more generally ^46,47^. In this framework, prior information provided by instruction or conditioning acts as a reference point (or *prior*, in Bayesian terminology) against which sensory input is evaluated. Priors play a pivotal role in shaping the perceptual process of an individual. Stronger, more precise priors exert greater influence on the underlying inferential process, shaping the perceived pain experience to align more closely with these expectations, even when the sensory evidence remains unchanged. This suggests that the effectiveness of a strategy lies in its ability to generate a robust prior, which in turn, leads to a more pronounced influence on subsequent perception. According to the Bayesian causal multi-source integration model, this depends not only on the quantity of prior information but also on the degree of agreement among information from various sources ^48^. When multiple sources provide consistent and complementary information (such as verbal instructions and experienced pain relief in the context of combined instructions and conditioning) and are inferred to share a common cause, these inputs are integrated during the inferential process and effectively increase the confidence in the prior. This, in turn, is likely to reduce variability across participants when instructions and conditioning are combined. On the contrary, the priors induced by instructions only – without providing any hands-on experience - are likely more driven by individual differences in personal history and varying subjective assumptions, explaining the higher variance we observed in behavioral placebo effects with INST, as compared to COND+INST.

Interestingly, our findings of stronger engagement of prefrontal regions by instructions paired with conditioning than instruction alone also align with observations from studies using dynamic reversal learning paradigms. In these paradigms, one strategy (e.g., conditioning) is used to induce learning and subsequent testing assesses whether another strategy (e.g., instructions) can reverse the learned effect. Studies employing this approach have demonstrated that the presence of instructions increases activation in the DLPFC ^18,21^, similarly to our study. Further, on the behavioral and peripheral-physiological level, instructions can modulate conditioned fear and autonomic responses (see ^18,19,49,50^), as well as brain responses associated with learning, including the VMPFC and striatum ^18,19,21^, two regions that were also found to mediate the effects of induction method on placebo analgesia our study. Interestingly, prior work suggests that the amygdala learns similarly regardless of instructions ^18,51^. Testing whether this finding extends to placebo analgesia would require studies that investigate effects of conditioning in the absence of instructions, which has not been tested with neuroimaging other than in the context of pain-predictive cues ^19^. This approach offers a promising way to explore the joint, dynamic impact of instructions and learning and may help to find the most robust and persistent way to induce placebo analgesia.

It could be argued that only some of the regions engaged during COND+INST are in fact causally related to the stronger analgesic effect. To test this assumption, we examined the correlation between brain responses and reported placebo analgesia across individuals. As expected, placebo responses in the brain were negatively associated with placebo analgesia across a widespread network of pain processing regions, including the thalamus, operculum, insula, middle cingulum and sensory regions. In contrast, no positive correlations survived correction for multiple comparisons across the whole brain.

Although there was no significant voxel-level interaction between brain-behavior correlation and induction type, we observed that the NPS, a weighted multivariate signature of pain ^13^ linked to nociception, predicted behavioral analgesia in the COND+INST condition more accurately than the stimulus intensity-independent pain signature (SIIPS; ^13^). This suggests that the COND+INST condition induces stronger top-down modulation of nociceptive processing, potentially explaining its enhanced analgesic effect. This is also in line with the outcome of our whole brain analysis showing stronger deactivation in nociceptive regions including the posterior insula and S1 in the condition combining instructions with conditioning.

Finally, we identified brain regions where the effect of induction type on behavioral placebo analgesia was mediated by brain placebo effects. The strongest ‘positive mediators’ were the VMPFC and the caudate, suggesting that these regions play a crucial role in translating the induction type into observable behavior. The VMPFC has long been associated with cognitive control ^52^, emotion regulation ^53^ and belief updating ^54^; furthermore, alterations in VMPFC activity and connectivity have been reported for different types of chronic pain ^55^.

The differential engagement of the VMPFC and caudate in placebo responses likely reflects the integration of richer, self-referential memories into expectations when conditioning is involved. The VMPFC shows increased activation during the retrieval of autobiographical memories, anchoring personal experiences within a narrative framework ^56^. Lesions in this region impair imagining detailed future scenarios, underscoring its role in weaving memories into prospective thinking ^54,57^, and lead to stronger reliance on instructions in the context of pain-predictive cues ^54^. This suggests, that in the context of placebo expectations, conditioning does more than reinforce expectancy – it likely integrates learned associations into a broader self-referential memory framework. Conditioning paired with instructions encourages participants to incorporate episodic experiences of the process (e.g., relief or positive outcomes) into their placebo expectations. This creates richer, more personally meaningful priors, in the Bayesian sense, that more strongly guide perception and behavior compared to abstract instructions alone Furthermore, the VMPFC is known to contain expected value signals that inform our choices ^58^. While INST, in itself, can establish a general prior belief about the expected effects of the (sham) treatment, only having hands-on experiences makes it possible to estimate the subjective “value” (efficacy) of the treatment. If, during the test phase, general expectations about the upcoming event (induced by verbal instruction) and concrete memories regarding the subjective value of the applied treatment (induced by conditioning) are inferred to be causally related, the two types of priors reinforce each other ^48^. This causal inference process will result in a stronger top-down modulatory force, consistent with the observed decreased activity in the ‘negative mediators’: the posterior insula, S2, and S1.

The caudate, part of the dorsal striatum, may complements this by integrating reward-related learning and supporting goal-directed behavior (e.g., ^59^). Conditioning may strengthen the role of the caudate in encoding the association between placebo cues and anticipated relief. Unlike the habit-focused putamen, the caudate facilitates goal-directed processes that align with purposeful expectancy of benefit. Together, the VMPFC and caudate, connected via corticostriatal circuits, may facilitate the integration of self-referential memory (VMPFC) and reinforcement history (caudate) into placebo expectations.

Our findings raise the question of whether the larger effect in the combined condition is simply due a stronger modulation of predictive processes (i.e., regardless of the technique or whether different types of learning are combined), due to the combination of different learning strategies (e.g., conditioning and verbal instruction), or due to the unique contribution of conditioning. In an early study, Colloca and colleagues demonstrated that increasing the number of conditioning sessions could enhance the placebo effect, implying that reinforcing learning through additional repetitions may strengthen the overall outcome ^24^. Schafer et al. (2015) ^25^ also showed that after several days of conditioning, placebo effects became larger and more resistant to changes in expectations; placebo effects persisted even when participants knew they had been taking a placebo and no longer expected relief. To investigate the influence of different strategies, van Lennep et al. (2023) ^60^ recently compared the effect of different combinations of conditioning, instruction, and observational learning on PA induction. Their results showed that the combination of all three strategies was superior to the use of any single strategy, but, surprisingly, it did not exceed the effectiveness of the combination of only two strategies – and interestingly, the combination of conditioning and instructions did not outperform conditioning alone.

It is important to highlight that our study did not specifically delineate the neural mechanisms of conditioning alone. While conditioning alone has been a prevalent method for modulating pain perception and pain-related fear in a plethora of studies ^61,62^, these investigations did not combine the conditioning procedure with treatment expectation. The results of our comparison between instructions and instructions combined with conditioning can therefore not necessarily be attributed to conditioning; instead, they are likely to be the result of an interaction between the two methods of inducing PA. Indeed, there are studies showing that conditioning alone, without suggestion or treatment expectations, shows limited efficacy in inducing PA ^63^.

In conclusion, our meta-analysis provides robust evidence for distinct neural correlates of PA induced by instructions alone and by a combination of instructions and conditioning. Our findings suggest that conditioning engages neural mechanisms beyond those of verbally induced PA. This improved understanding of the efficacy and underlying mechanisms of different placebo induction techniques not only informs ways to maximize and exploit placebo effects in experimental studies. It has the potential to guide targeted interventions in clinical settings to optimize analgesic strategies, including for those with an impaired ability to amplify analgesic outcome through placebo mechanisms, such as patients with neurodegenerative diseases ^64,65^.

## Materials and Methods

### Data acquisition

The present study is a systematic meta-analysis of individual participant data across *k* = 16 peer-reviewed, within-participant placebo neuroimaging studies (see Table 1 for an overview of all included studies). Data from the included studies, together with data from four more between-participant studies, served as a basis for previous meta-analyses, examining placebo effects on the NPS ^14^ and across the entire brain ^15^. In contrast to these previous analyses, the scope of the present analysis was restricted to within-participant studies, allowing us to consider the individual magnitude of PA when investigating the neural correlates of PA induced by verbal instructions alone (INST) or by verbal instructions together with conditioning (COND-INST). Our analysis included single-participant data from a total of *n* = 415 participants (*n* = 409 after excluding participants with missing pain ratings).

A detailed description of the data collection procedures can be found in ^14^. In brief, we first performed a systematic literature search to identify experimental functional magnetic resonance imaging (fMRI) studies of PA. Eligible studies met the following criteria: (a) published in English in a peer-reviewed journal; (b) constituted an empirical investigation; (c) involved human participants; (d) employed functional neuroimaging of the brain during evoked pain; and (e) induced pain under matched placebo and control conditions. For the present study, we added that (f) studies had to be conducted within-participant, with each participant serving as their own control and (g) availability of pain ratings in both the placebo and the control conditions. Authors of all identified studies were contacted and invited to share their data. We collected single-participant, first-level, whole-brain standard space summary images of pain response (statistical parametric maps) from the original analyses, corresponding pain ratings separately for placebo and control conditions, experimental design parameters, and demographic data. From studies that applied conditioning, we only used data from the test phase (and not the conditioning phase). The included studies induced PA either by verbal instructions alone (INST; *k* = 5, *n* = 147) or by a combination of verbal instructions and conditioning (COND-INST; *k* = 11, *n* = 268, see Table 1). Seven studies used a topical cream as a placebo, five used an intravenous infusion, and two used sham acupuncture. One study each used either sham transcutaneous nerve stimulation (TENS) or a placebo nasal spray. The majority of studies employed thermal stimulation (*k* = 11), followed by laser or distension (each *k* = 2) and electrical pain (*k* = 1; see **Supplementary Figures S1-S3** for details). Details on the conditioning procedures used in the different studies can be found in **Supplementary Table S4**. Detailed risk of bias reports can be found in Zunhammer et al. (2021).

### Harmonization

#### Behavioral Data

To construct harmonized measures of the behavioral placebo response across studies, we first converted pain intensity ratings to a score between 0-100 and then subtracted the placebo condition ratings from the control condition ratings (positive values correspond to analgesia). Additionally, we obtained study-wise standardized effect size summaries of the behavioral PA by calculating Hedges’ g scores. Hedges’ *g*, as is common in meta-analyses ^66^. Similarly to the Cohen *d*, Hedges *g* is based on the mean difference between conditions divided by standard deviation, but with an additional correction for small sample bias. Specifically, as only within-participant studies were included, we used Hedges *g*rm, which is based on the SD of within-participant differences corrected for within-participant correlations (for details see ^14,15^).

#### Neuroimaging Data

To address center effects and differences in image acquisition procedures across studies, we harmonized the brain data (first-level control-placebo contrast images) using quantile transformation, applied independently to each voxel within each study. Quantile transformation, similar to ranking, is a widely used data harmonization technique ^66^ that assigns each data point its quantile within the study, resulting in a value between 0 and 1. This process forces the transformed data to follow a uniform distribution, thereby eliminating differences in data distributions across studies. However, the conventional form of quantile transformation also removes all information about study-specific means, making it unsuitable for comparing effects across studies (e.g., assessing the overall effects of induction techniques on PA, as in path A of a mediation analysis). This limitation can be overcome by using a zero-preserving quantile transformation, which involves converting the original values to their quantile distance from zero. Specifically, we calculated:

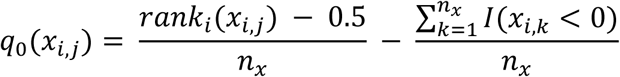

where *q*_0_ is the zero-preserving quantile transformation, *x*_*i*,*j*_ is the value in voxel a given voxel in participant j from stud i, *rank*_*i*_(.) denotes the rank transformation across all participants in study i, using average ranking for ties, and *I*(*x*_*i*,*k*_ < 0) is an indicator function that equals 1 if *x*_*i*,*k*_ is positive and 0 otherwise. This equation describes how each voxel value in the individual level placebo brain images is transformed by ranking it within its study, normalizing the rank (to get quantiles), and adjusting for the proportion of negative values in the batch.

### Data analysis

#### Standardized Behavioral Summary Data

The effect of induction technique on the standardized effect size summaries of behavioral PA across studies was investigated by a *T*-test comparing the Hedges’ *g*-values between the two groups of studies.

#### Mediation Analysis

To investigate the neural correlates of the effect of induction type on behavioral PA, we performed a voxel-wise mediation analysis, with the main aim of evaluating the indirect effects of induction type (X) on behavioral placebo analgesia (Y) through placebo-related neural activity as a mediator variable (*M*). Throughout our mediation analysis, we corrected for age, sex and control pain rating, and modelled studies with dummy regressors orthogonalized to induction type. In detail, our approach consisted of fitting three different ordinary least squares linear models with a costume analysis code written in python (see section on Code Availability).

First, the mediator M (brain data) is modeled as a function of the independent variable X (induction type) and covariates using linear regression. This step estimates the effect of induction type on brain activity (*path a*). Second, the outcome Y (behavioral PA) is modeled as a function of the mediator M (brain activity) and covariates using linear regression. This step estimates the correlation of brain activity changes with behavioral placebo analgesia across participants (*path b*). Finally, the indirect effect (also known as the average causal mediation effect or ACME) is calculated as the product of the coefficients from *path a* and *path b*.

Furthermore, we estimated the group-mean brain activity, assuming dummy coding for the two groups of studies. The reference group mean is derived from the intercept, and the other group’s mean is adjusted by the group difference.

The significance of the path a and path b effects, the indirect (mediation) effect and the group-mean activations were assessed by repeating the whole analysis on 10000 surrogate datasets, each constructed by flipping the sign of the brain data for randomly selected participants and rerunning the whole procedure, including harmonization and mediation analysis. Note that the approach is equivalent to permutation testing on with considering the two brain maps (placebo and control pain) of a participant as an exchangeability group.

The actual (unflipped) effect estimates were contrasted to the resulting null-distribution to obtain uncorrected p-values which were adjusted for multiple comparisons across the brain by false discovery rate (FDR) correction. As the precision of the estimation of very low p-values with permutation testing is limited by the number of permutations, we used tail approximation to calculate more accurate p-values ^67^. Specifically, the sign flipping-based null-distribution was subject of tail approximation (in each voxel independently), by fitting a generalized Pareto distribution to the tail of the distribution ^68^. The optimal tail ratio was selected from a pre-defined set of tail rations (from 0.1 to 0.3, with increments of 0.05), based in the goodness of fit, as measured by the Kolmogorov-Smirnov statistic ^69^. P-values were then approximated based on the fitted Pareto distribution:

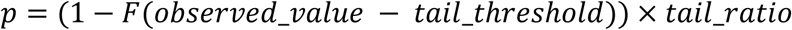

Where *F*(x) is the Cumulative Distribution Function of the generalized Pareto distribution.

### CA1: Variance analysis

We investigated whether the variance of the behavioral placebo response (unharmonized but converted to VAS 0-100) differs between INST and COND+INST in a two-step approach. First, we fitted a mixed effect model that explains behavioral analgesia with age, sex and control pain rating (all centered) as fixed effects and study as random effect. We then performed Levene’s test of equal variances on the residuals of this model, to compare the variance of the residuals in the INST vs. COND+INST studies.

### CA2: Conjunction Analysis

We performed a conjunction analysis on the two group-mean maps (mean placebo brain response to INST and COND+INST) by taking the maximum p-value from the two images in each voxel, and correcting for false discovery rate across brain voxels, as described in ^70^.

### CA3: Habenula ROI analysis

We obtained the Habenula ROI from the subcortical brain atlas of ^71^, divided it into bilateral components (by x = 0). Within all voxels of each ROI, we averaged the voxel-wise path a, path b and mediation (a*b) regression coefficients, as well as the group-mean estimates for INST and COND+INST (see ‘Mediation Analysis’). To construct the null distribution for statistical inference, we used the ROI-wide averages of the voxel-wise null-distribution estimates, calculated based on sign-flipped 10000 surrogate datasets, separately for path a, b, a*b and the INST and COND+INST group means. Similarly to the voxel-wise analysis, we calculated p-values by fitting a generalized Pareto distribution on the tail of the null distributions.

### CA4: Pain signatures

To test whether placebo analgesia induced by the two techniques scaled with changes in the overall activity in pain-related brain regions, we correlated the placebo analgesic effect with activation changes in two established neural signatures of pain, namely the Neurologic Pain Signature (NPS; ^13^) and the Stimulus Intensity Independent Pain Signature (SIIPS; ^26^), separately for the COND-INST and INST. Engagement of the NPS was been shown to robustly correlate with pain induced by different levels of noxious input ^13,14,72^, whereas activity in the SIIPS reflects stimulus-independent variations in reported pain intensity ^26^. When comparing the outputs of the investigated neural signatures (NPS and SIIPS), we first computed the output of the signatures for each participant as the dot product of the signatures’ predictive coefficient map and the participants’ harmonized brain imaging data. The resulting signature scores were then fed into a linear model, *signature response ∼ induction type * behavioral PA + CTR pain rating + age + sex + study*. As in the mediation analysis, studies were modelled with dummy regressors orthogonalized to induction type. For all coefficients of interest, we performed bootstrapping (with 10000 bootstrap samples) to obtain 95% confidence intervals and p-values.

### Visualization

MRIcroGL (v28.5.2017) and the python package *nilearn* was used to create illustrations of statistical parametric maps. Activation peaks were summarized using the Mars brain atlas and the Yeo 7-network functional parcellation, based on the MNI-coordinates of the peak. All result maps follow the neurological convention, where the left side corresponds to left hemisphere in coronal and axial sections.

On **Figures 3-5**, when visualizing uncorrected results (*p* < .01), we only show clusters that had at least one voxel that survived false discovery rate correction (*q* < 0.05).

Activation peaks were summarized using the Mars brain atlas ^40^ and the Yeo 7-network functional parcellation ^41^, based on the MNI-coordinates of the peak.

## Data availability

Participant-level source data are available from the authors upon reasonable request and with permission from the Placebo Imaging Consortium.

## Code availability

Analysis code for carrying out data cleaning is available at: https://github.com/mzunhammer/PlaceboImagingMetaAnalysis.

The complete code for reproducing the analyses and figures presented in the manuscript are available at: https://github.com/pni-lab/placebo-conditioning-meta-analysis.

## Contributions

U.B. and T.D.W. had full access to all the data in the study and take responsibility for the integrity of the data and the accuracy of the data analysis. Concept and design: T.S., U.B., and T.D.W. Acquisition of the data: All authors. Drafting of the manuscript: T.S., H.H., K.W., T.W., U.B. Critical revision of the manuscript for important intellectual content: All authors. Statistical analysis: B.K., T.S. Visualization of Results: T.S., B.K., H.H., Obtained funding: U.B., T.S. Administrative, technical, or material support: U.B. Supervision: U.B. and T.D.W.

## Acknowledgements

This research was funded by the German Research Foundation, TRR 289 Treatment Expectation (Gefördert durch die Deutsche Forschungsgemeinschaft (DFG) – Projektnummer 422744262 – TRR 289; U.B., T.S., H.H., K.W.) and the National Institutes of Health R37 MH076136 (T.D.W.).

