## Supplementary Materials for "Meta-analytic evidence for distinct neural correlates of conditioned vs. verbally induced placebo analgesia"

for the manuscript entitled

by

Tamas Spisak, Helena Hartmann, Matthias Zunhammer, Balint Kincses,
Katja Wiech, Tor D. Wager, & Ulrike Bingel, for the Placebo Imaging Consortium

### Supplementary figures

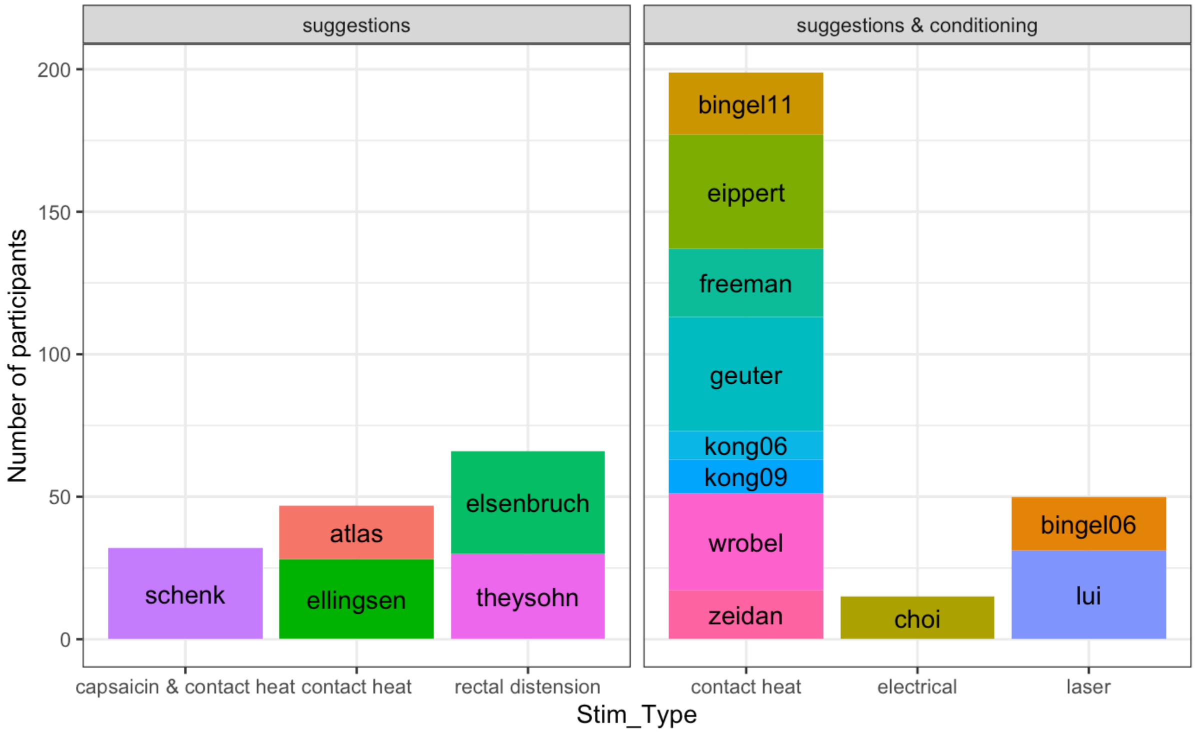


*Supplementary Figure S1. Number of participants and studies grouped by the type of pain stimulation.*

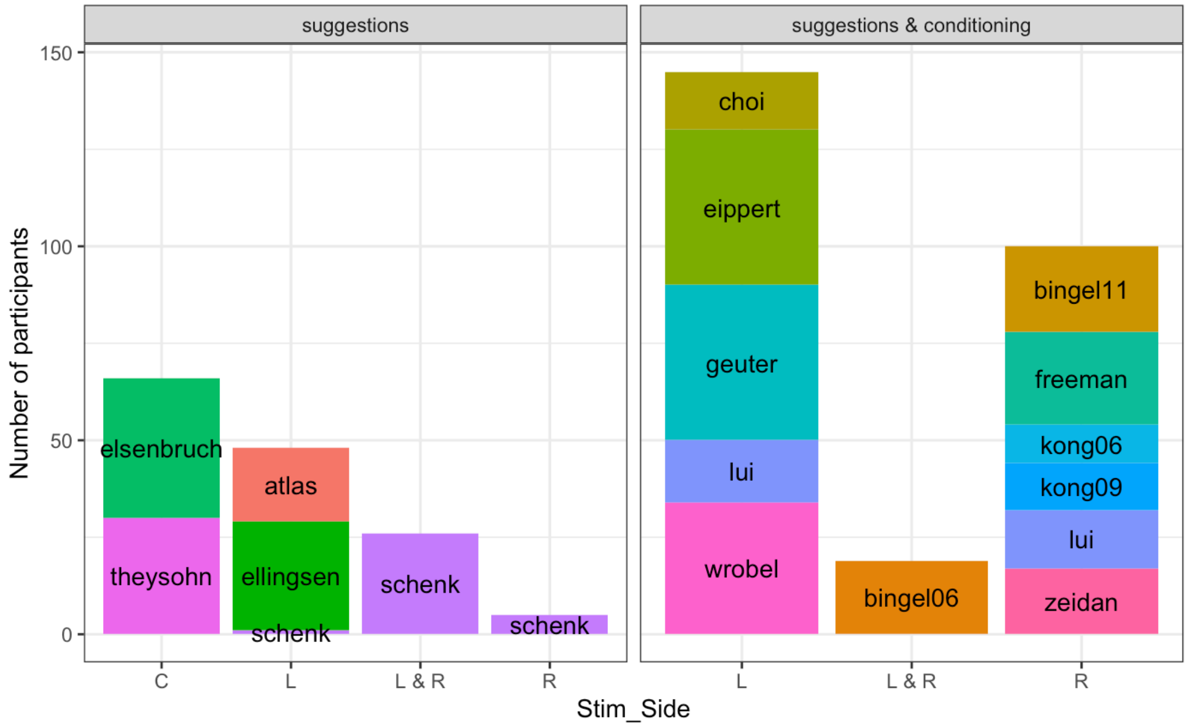


*Supplementary Figure S2. Number of participants and studies grouped by the laterality of pain stimulation. L: left, R: right, C: no laterality (rectal distension studies).*

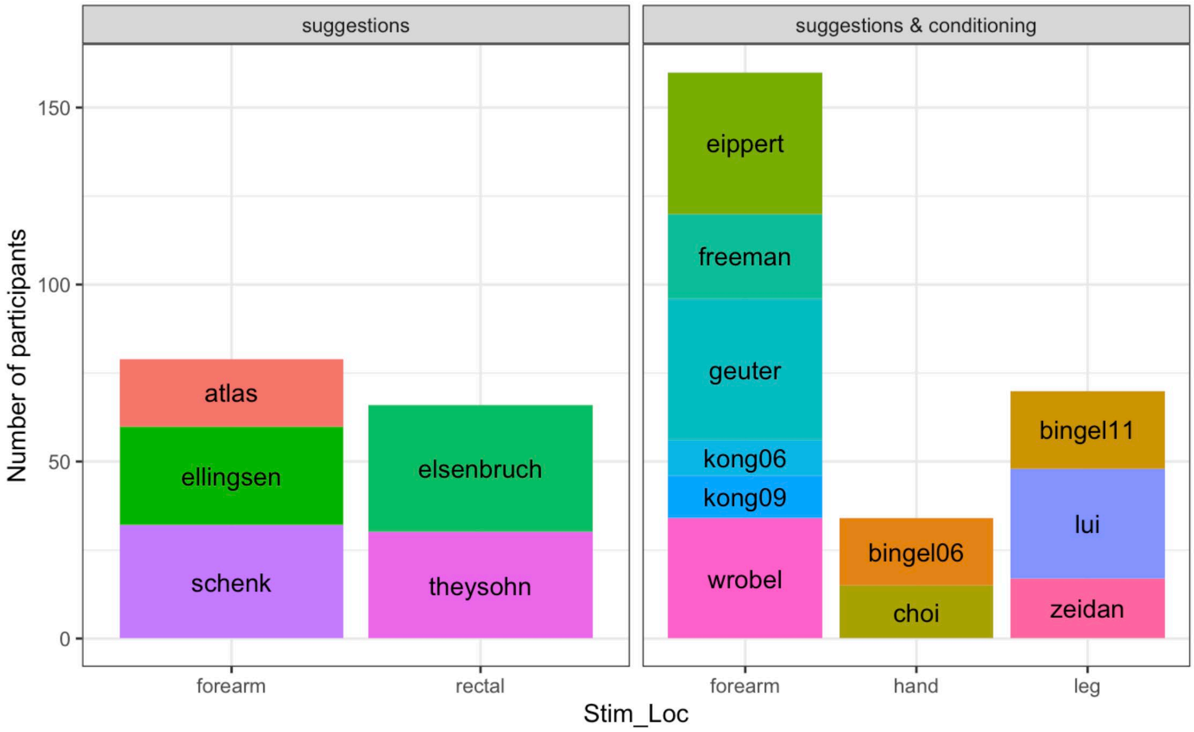


*Supplementary Figure S3. Number of participants and studies grouped by the location of pain stimulation.*

### Supplementary tables

*Supplementary Table S1. Results of the random effect model that explains placebo rating differences with age, sex and control pain rating as fixed effects and study as random effect. The residuals of this model were subject to Levene's test of equal variances to compare INST vs. COND+INST studies.*

| | ***coef*** | ***std err*** | ***z*** | ***p(\|T\|>0)*** | ***95% CI (coef)*** | |
|---|---|---|---|---|---|---|
| *intercept* | 11.052 | 1.267 | 8.726 | **< 0.0001** | 8.570 | 13.535 |
| *sex* | 2.310 | 1.832 | 1.261 | 0.207 | -1.281 | 5.901 |
| *age* | -0.301 | 0.140 | -2.146 | **0.032** | -0.576 | -0.026 |
| *control pain rating* | 0.273 | 0.041 | 6.728 | **< 0.0001** | 0.193 | 0.352 |
| *study* | 15.044 | 0.653 | | | | |

*Supplementary Table S2. Full table with regression coefficients, standard errors, T-scores, p-values and confidence intervals for the statistical test modelling the effect of induction type on behavioral placebo analgesia.*

| | ***coef*** | ***std err*** | ***T*** | ***p(\|T\|>0)*** | ***p(T>0)*** | ***95% CI (coef)*** | |
|---|---|---|---|---|---|---|---|
| *intercept* | 11.1459 | 0.774 | 14.394 | | **< 0.0001** | 9.623 | 12.668 |
| *placebo induction* | 1.3394 | 0.807 | 1.660 | | **0.049** | -0.247 | 2.926 |
| *control pain rating* | 0.3049 | 0.041 | 7.404 | **< 0.0001** | | 0.224 | 0.386 |
| *age* | -0.2898 | 0.146 | -1.981 | **0.048** | | -0.577 | -0.002 |
| *sex* | 2.2475 | 1.920 | 1.170 | 0.243 | | -1.528 | 6.023 |
| *Atlas* | -0.1930 | 0.677 | -0.285 | 0.776 | | -1.523 | 1.137 |
| *Bingel 06* | 2.1391 | 0.721 | 2.966 | **0.003** | | 0.721 | 3.557 |
| *Bingel 11* | 0.8278 | 0.713 | 1.161 | 0.246 | | -0.574 | 2.230 |
| *Choi* | -0.7059 | 0.738 | -0.957 | 0.339 | | -2.157 | 0.745 |
| *Eippert* | 0.3153 | 0.686 | 0.460 | 0.646 | | -1.033 | 1.664 |
| *Ellingsen* | 0.5167 | 0.635 | 0.813 | 0.416 | | -0.732 | 1.766 |
| *Elsenbruch* | 0.2861 | 0.598 | 0.479 | 0.632 | | -0.889 | 1.461 |
| *Freeman* | 0.5094 | 0.710 | 0.717 | 0.474 | | -0.887 | 1.906 |
| *Geuter* | -1.7419 | 0.690 | -2.525 | **0.012** | | -3.098 | -0.385 |
| *Kong 06* | -1.7226 | 0.754 | -2.284 | **0.023** | | -3.206 | -0.240 |
| *Kong 09* | -1.4959 | 0.752 | -1.989 | **0.047** | | -2.974 | -0.018 |
| *Lui* | 1.2096 | 0.766 | 1.579 | 0.115 | | -0.297 | 2.716 |
| *Schenk* | -1.6031 | 0.611 | -2.625 | **0.009** | | -2.804 | -0.402 |
| *Theysohn* | 0.9933 | 0.667 | 1.489 | 0.137 | | -0.318 | 2.305 |
| *Wrobel* | 0.1456 | 0.683 | 0.213 | 0.831 | | -1.196 | 1.488 |
| *Zeidan* | -0.2074 | 0.734 | -0.282 | 0.778 | | -1.651 | 1.236 |

*Supplementary Table S3. Full table including the regression coefficients, standard errors, T-scores, p-values and confidence intervals for the statistical test modelling the interaction effect between induction type and behavioral analgesia on NPS scores.*

| | ***coef*** | ***std err*** | ***T*** | ***p*** | ***95% CI (coef)*** | |
|---|---|---|---|---|---|---|
| ***intercept*** | 0.0011 | 0.002 | 0.670 | 0.72 | -0.0026 | 0.0047 |
| ***placebo induction*** | 5.17e-05 | 0.002 | 0.028 | 0.52 | -0.0037 | 0.00386 |
| ***analgesia*** | -0.0004 | 0.001 | -3.559 | **0.0005** | -0.0006 | -0.00016 |
| ***analgesia:induction*** | -0.0002 | 8.89e-05 | -1.825 | **0.034** | -0.000344 | -0.000001 |
| **control pain rating** | -6.414e-05 | 8.42e-05 | -0.762 | 0.21 | -0.0002 | 0.0001 |
| ***age*** | -0.0006 | 0.001 | -2.060 | **0.014** | -0.001 | -0.0001 |
| ***sex*** | 0.0001 | 0.004 | 0.040 | 0.51 | -0.0069 | 0.0072 |
| ***Atlas*** | 0.0007 | 0.001 | 0.566 | 0.572 | -0.002 | 0.003 |
| ***Bingel 06*** | -0.0013 | 0.001 | -0.918 | 0.359 | -0.004 | 0.001 |
| ***Bingel 11*** | 0.0011 | 0.001 | 0.808 | 0.420 | -0.002 | 0.004 |
| ***Choi*** | -0.0018 | 0.001 | -1.259 | 0.209 | -0.005 | 0.001 |
| ***Eippert*** | 0.0010 | 0.001 | 0.791 | 0.429 | -0.002 | 0.004 |
| ***Ellingsen*** | 0.0004 | 0.001 | 0.360 | 0.719 | -0.002 | 0.003 |
| ***Elsenbruch*** | -0.0014 | 0.001 | -1.215 | 0.225 | -0.004 | 0.001 |
| ***Freeman*** | 0.0034 | 0.001 | 2.499 | **0.013** | 0.001 | 0.006 |
| ***Geuter*** | 0.0021 | 0.001 | 1.566 | 0.118 | -0.001 | 0.005 |

| | | | | | | |
|---|---|---|---|---|---|---|
| ***Kong 06*** | 0.0012 | 0.001 | 0.856 | 0.393 | -0.002 | 0.004 |
| ***Kong 09*** | -0.0011 | 0.001 | -0.767 | 0.444 | -0.004 | 0.002 |
| ***Lui*** | -0.0003 | 0.001 | -0.209 | 0.834 | -0.003 | 0.003 |
| ***Schenk*** | -0.0021 | 0.001 | -1.753 | 0.080 | -0.004 | 0.000 |
| ***Theysohn*** | 0.0026 | 0.001 | 2.006 | **0.046** | 5.11e-05 | 0.005 |
| ***Wrobel*** | -0.0028 | 0.001 | -2.126 | **0.034** | -0.005 | -0.000 |
| ***Zeidan*** | -0.0026 | 0.001 | -1.836 | 0.067 | -0.005 | 0.000 |

*Supplementary Table S4. Full table including the regression coefficients, standard errors, T-scores, p-values and confidence intervals for the statistical test modelling the interaction effect between induction type and behavioral analgesia on SIIPS scores.*

| | ***coef*** | ***std err*** | ***T*** | ***p*** | ***95% CI (coef)*** | |
|---|---|---|---|---|---|---|
| ***intercept*** | 0.0063 | 0.004 | -1.455 | 0.074 | -0.015 | 0.0023 |
| ***placebo induction*** | -0.0017 | 0.004 | -0.419 | 0.359 | -0.010 | 0.0065 |
| ***analgesia*** | -0.0008 | 0.001 | -3.208 | **0.004** | -0.0013 | -0.00019 |
| ***analgesia:induction*** | -0.0002 | 0.001 | -1.168 | 0.145 | -0.0007 | 0.0002 |
| **control pain rating** | -9.871e-05 | 0.001 | -0.523 | 0.28 | -0.0005 | 0.0003 |
| ***age*** | -8.955e-05 | 0.001 | -0.142 | 0.44 | -0.0011 | 0.0013 |
| ***sex*** | -0.0032 | 0.008 | -0.386 | 0.34 | -0.019 | 0.013 |
| ***Atlas*** | -0.0016 | 0.003 | -0.547 | 0.585 | -0.007 | 0.004 |
| ***Bingel 06*** | -0.0039 | 0.003 | -1.218 | 0.224 | -0.010 | 0.002 |
| ***Bingel 11*** | 0.0042 | 0.003 | 1.363 | 0.174 | -0.002 | 0.010 |
| ***Choi*** | 0.0034 | 0.003 | 1.089 | 0.277 | -0.003 | 0.010 |
| ***Eippert*** | -0.0015 | 0.003 | -0.520 | 0.603 | -0.007 | 0.004 |
| ***Ellingsen*** | -0.0014 | 0.003 | -0.512 | 0.609 | -0.007 | 0.004 |
| ***Elsenbruch*** | 0.0012 | 0.003 | 0.455 | 0.649 | -0.004 | 0.006 |
| ***Freeman*** | 0.0043 | 0.003 | 1.419 | 0.157 | -0.002 | 0.010 |
| ***Geuter*** | 0.0059 | 0.003 | 1.971 | 0.049 | 1.32e-05 | 0.012 |
| ***Kong 06*** | -0.0060 | 0.003 | -1.836 | 0.067 | -0.012 | 0.000 |
| ***Kong 09*** | -0.0032 | 0.003 | -0.974 | 0.331 | -0.010 | 0.003 |
| ***Lui*** | 0.0009 | 0.003 | 0.258 | 0.796 | -0.006 | 0.007 |
| ***Schenk*** | -0.0015 | 0.003 | -0.575 | 0.565 | -0.007 | 0.004 |
| ***Theysohn*** | 0.0030 | 0.003 | 1.045 | 0.297 | -0.003 | 0.009 |
| ***Wrobel*** | -0.0051 | 0.003 | -1.736 | 0.083 | -0.011 | 0.001 |
| ***Zeidan*** | -0.0022 | 0.003 | -0.686 | 0.493 | -0.008 | 0.004 |

*Supplementary Table S5. Types of conditioning separate for each included study.*

| **Study** | **Induction type** | **Treatment** | **Cond type** | **Cond session** | **Cond length** |
|---|---|---|---|---|---|
| Atlas et al. (2012) | | IV-infusion | NA | NA | NA |
| Bingel et al. (2006) | V + C | Topical cream | Response | Same session | 2x4 stimuli per hand |
| Bingel et al. (2011) | V + C | IV-infusion | Response | Extra session[a] | No info |
| Choi et al. (2011) | V + C | IV-infusion | Response | Same session | No info |
| Eippert et al. (2009) | V + C | Topical cream | Response | Extra & same session[b] | 6 stimuli per session |
| Ellingsen et al. (2013) | V | Nasal spray | NA | NA | NA |
| Elsenbruch et al. (2012) | V | IV-infusion | NA | NA | NA |
| Freeman et al. (2015) | V + C | Topical cream | Response | Extra session[c] | 3 stimuli |
| Geuter et al. (2013) | V + C | Topical cream | Response | Same session[d] | 12 stimuli[e] |
| Kong et al. (2006) | V + C | Sham acupuncture | Response | Extra & same session[f] | 6/12 stimuli per session |
| Kong et al. (2009) | V + C | Sham acupuncture | Response | Extra & same session[g] | 8/6 stimuli per session |
| Lui et al. (2010) | V + C | Sham TENS | Response | Same session | 2x12 stimuli |
| Schenk et al. (2014) | V | Topical cream | NA | NA | NA |
| Theysohn et al. (2014) | V | IV-infusion | NA | NA | NA |
| Wrobel et al. (2014) | V + C | Topical cream | Response | Extra & same session[h] | 12/6 stimuli per session |
| Zeidan et al. (2015) | V + C | Topical cream | Response | Extra sessions[i] | 10 stimuli per session |

*Note*. IV = intra-venous; TENS = transcutaneous electrical nerve stimulation; V = verbal; C = conditioning; NA = not applicable; Response = response conditioning where pain is lowered in placebo condition. [a] Cond session min. 24hrs before test session. [b] Cond on consecutive Day 1 in the lab and Day 2 in the scanner before test session. [c] Sessions separated by 2-14 days. [d] Two sessions around one week apart for weak and strong placebo. [e] Six stimuli each outside and inside the scanner. [f] Sessions min. four days apart. [g] Sessions min. four days apart. [h] Cond on consecutive Day 1 and Day 2 outside of the scanner. [l] Four-day placebo conditioning.
